# Epipelagic to mesopelagic variability of acoustic backscatter in the California Current

**DOI:** 10.64898/2026.04.14.718518

**Authors:** J. Guiet, C. Wall, K. Srinivasan, D. Bianchi

## Abstract

Mid-Trophic Level (MTL) organisms—including krill, forage fish, and mesopelagic fish— are abundant in the California Current System (CCS) and play an essential role in transferring energy and biomass from primary producers to top predators. However, their spatiotemporal distribution and variability remain poorly understood, particularly with respect to vertical structure across epipelagic and mesopelagic habitats and coastal-offshore gradients. This lack of understanding emerges from both the complexity of MTL interactions with a heterogeneous environment and the challenges associated with sampling these organisms at high spatial and temporal resolution. To address this gap, we analyze 11 years of fisheries acoustic observations in the CCS (2006-2016) to characterize the spatiotemporal dynamics of MTLs as inferred from acoustic backscatter. Acoustic observations at 38 and 120 kHz, collected during day and night across depth strata from 15 to 495 m, reveal consistent cross-shore, seasonal, and latitudinal patterns in the backscatter of acoustically defined zooplankton, epipelagic fish, and mesopelagic fish communities. These patterns include: (1) weaker cross-shore gradients in mesopelagic relative to epipelagic communities; (2) a temporal succession among communities associated with seasonal upwelling; and (3) a multimodal latitudinal distribution with distinct coastal backscatter peaks. We further investigate relationships between acoustic backscatter and co-located environmental variables from in situ, remote sensing, and reanalysis products to elucidate plausible mechanisms underlying MTL dynamics.

**Highlights:**

- Fisheries acoustics resolve variability in mid-trophic communities
- Eleven years of backscatter reveal consistent patterns in the California Current
- Epipelagic backscatter declines faster from the coast to offshore than mesopelagic
- Seasonal changes in community composition are linked to upwelling dynamics
- Backscatter exhibits multimodal latitudinal distributions with distinct peaks

## 1 Introduction

Mid-trophic levels (MTL), including macrozooplankton, forage fish, and mesopelagic fish, are fundamental components of marine ecosystems that mediate the transfer of biomass and energy from plankton to large predators [1, 2, 3]. MTLs affect ocean biogeochemistry via top-down grazing on primary producers [4], by generating rapidly sinking particles and modulating particle transformations [5, 6], and through diel vertical migrations [7, 8, 9]. MTLs comprise a variety of taxa and species, ranging from near-surface forage fish such as sardines and anchovies, to invertebrates including squid and krill, to vertically migrating and resident mesopelagic fish such as myctophids and bristlemouths [10, 11, 12].

Because of their ecological role and economic value, MTL are routinely surveyed in support of fisheries assessments, including in the California Current System (CCS) [13]. In this productive eastern boundary upwelling system, numerous studies have documented variability in MTL across a range of spatial and temporal scales. These include responses to episodic wind-driven upwelling and associated fronts, filaments, and eddies that generate environmental heterogeneity [14, 15, 16]; seasonal shifts in the spatial distributions of MTL assemblages [17, 18, 19]; and interannual changes in community abundance [20] and composition [10, 11].

In the CCS, the timing, location, and intensity of seasonal upwelling are dominant drivers of variability [21]. Primary production and phytoplankton biomass typically peak early in the upwelling season and nearshore, where coastal upwelling injects nutrients into the euphotic zone [21, 22, 23, 24]. In contrast, zooplankton biomass tends to peak progressively later in the season and increasingly offshore or downstream of upwelling centers [25, 26, 27]. This spatial and temporal trophic succession is thought to arise from a combination of offshore advection by Ekman transport and geostrophic surface currents, trophic-level-dependent growth rates that delay biomass accumulation at higher trophic levels, and biomass aggregation and retention within mesoscale eddies and along frontal structures [28, 29, 27, 30, 31]. This schematic view provides a first-order description of trophic succession in the CCS. In practice, biological responses, particularly at higher trophic levels, are often spatially and temporally heterogeneous, reflecting variability in physical and ecological drivers as well as observational limitations [21, 23]. The extent to which this framework, typically applied to lower trophic levels and zooplankton, extends to MTL fish remains unclear.

Although stronger upwelling is associated with enhanced primary production [32], its impacts on MTL species and fish communities are often mixed and less predictable. For instance, shifts between strong and weak upwelling regimes tend to favor anchovies or sardines, respectively, because of morphological differences that affect feeding behavior and prey preference [33]. Variations in upwelling intensity can also compress and expand pelagic habitats, alternately favoring offshore versus coastal assemblages dominated, respectively, by coastal pelagic, mesopelagic and subtropical species, and by young-of-the-year groundfish and krill [34, 35, 36]. Upwelling modulates the occurrence of hypoxia at depth, thereby influencing the distribution and composition of mesopelagic communities [37, 38]. Beyond upwelling, currents also affect MTL by redistributing water masses and their resident organisms. For example, interannual changes in transport by the California Current and subsurface Undercurrent are linked to shifts between northern (transition-zone/sub-Arctic) and southern (tropical/subtropical) fauna [2, 39]. At local scales, currents generate retention zones that support biological hotspots [40, 27], transport communities and favorable conditions via eddies [41], aggregate or disperse organisms [14, 42], and form fronts that can act as barriers between communities [43, 15].

While this range of processes makes it difficult to characterize MTL variability at the ecosystem scale, observations of synchronous community-level responses point to an important role for large-scale climatic influences [11]. For instance, the abundance and distribution of epipelagic organisms is affected by shifts between “warm” (weakened upwelling) and “cold” (enhanced upwelling) phases associated with El Niño-Southern Oscillation (ENSO), the Pacific Decadal Oscillation (PDO) and the North Pacific Gyre Oscillation (NPGO) [44, 45]. Synchronous responses to climatic forcings, such as favorable upwelling conditions, propagate through the water column—from surface to mid-water and benthic communities—altering biomass, albeit with a damped response linked to different species longevity [30]. Multi-annual events such as the 2015–2016 marine heatwave in central California have been observed to uniformly shift the vertical distribution of mesopelagic MTLs [38]. On longer timescales, the northward inflow of water masses of tropical origin has increased the presence of warm-water-associated mesopelagic species in Southern California [10]. As evidence increasingly sheds light on the community-level dynamics of MTLs in the CCS, connections between epipelagic and mesopelagic communities remain less clearly resolved—particularly at seasonal timescales—despite their central role in pelagic food webs [46, 47].

In this study, we build on more than a decade of active fisheries acoustic observations in the CCS [48, 49, 13, 50] to analyze variability in MTLs from the epipelagic to mesopelagic zones, spanning cross-shore and meridional gradients. Multi-frequency echosounder observations provide a high-resolution view of the distribution of MTLs for dominant functional types [51, 52, 53]. We merge a total of 41 surveys (Table 1) to examine epipelagic-mesopelagic linkages by analyzing variations in acoustic backscatter—a proxy for presence, abundance, and community composition—together with two indicators of vertical structure: center of mass and vertical spread. We characterize these patterns along three well-sampled spatiotemporal dimensions: cross-shore and seasonal variability in Southern California, and a coast-wide alongshore gradient during peak upwelling. Finally, we extend the conceptual framework of trophic succession to MTL communities and relate observed patterns to environmental variability, shedding light on mechanisms underlying MTL dynamics.

**Table 1:** List of acoustic transects in the CCS from the following programs: Integrated Acoustic Survey of Pacific Hake (HAKE); Integrated Acoustic Survey of Pacific Hake and Pacific Sardines (SAKE); California Current Ecosystem Survey (CCE); CalCOFI; Coastal Pelagic Species (CPS). Calibration status denotes transects with available calibration parameters, applied or updated during processing (C); transects calibrated using indirect calibration information (C^∗^); and transects processed with default, pre-recorded calibration parameters lacking additional documentation (D).

| Ship | Cruise | Program | Year | Dates (nb. days) | 18 | 38 | 70 | 120 | 200 | Sensor depth | Range | Roll/Pitch | Calibration | DOI (https://doi.org/) |
| --- | --- | --- | --- | --- | --- | --- | --- | --- | --- | --- | --- | --- | --- | --- |
| Miller Freeman | MF0706 | CCE | 2007 | 06/05-06/10 (6) | X | X | - | X | X | 9.1 | 0-500 | - | C* | 10.25921/rnmq-wg11 |
|  | MF0710 | HAKE | 2007 | 06/20-08/21 (54) | X | X | - | X | X | 9.1 | 0-750 | - | C* | 10.7289/V5Q81B1B |
|  | MF1002(1004) | CalCOFI | 2010 | 04/26-05/17 (17) | X | X | - | X | X | 9.1 | 0-650 | - | C* | 10.25921/042e-pj65 |
| Oscar Dyson | DY0604 | CCE | 2006 | 04/06-05/08 (31) | X | X | X | X | X | 7.4 | 0-500 | - | D | 10.25921/y2rj-0g29 |
| David Starr Jordan | DS0604 | CCE | 2006 | 04/06-04/21 (12) | - | X | X | X | X | 3.75 | 0-500 | - | C | 10.25921/tzx1-zc28 |
|  | DS0704 | CalCOFI | 2007 | 03/27-05/01 (36) | - | X | X | X | X | 3.75 | 0-500 | - | C* | 10.25921/t8wz-3w11 |
| Ocean Starr | OS1207 | CalCOFI | 2012 | 07/10-07/27 (18) | - | X | X | X | X | 3.75 | 0-750 | - | C | 10.25921/v64z-vr54 |
|  | CCE1207 | CCE | 2012 | 07/28-08/25 (29) | - | X | X | X | X | 3.75 | 0-1000 | - | C | 10.6075/J0TX3CPV |
|  | OS1403 | CalCOFI | 2014 | 04/03-04/30 (24) | - | X | X | X | X | 3.75 | 0-750 | - | C* | 10.25921/2s4s-0579 |
| New Horizon | NH0901 | CalCOFI | 2009 | 01/07-01/24 (18) | X | X | X | X | X | 3.75 | 0-750 | - | C | 10.25921/rw77-wb44 |
|  | CCE0904 | CCE | 2009 | 04/22-05/03 (12) | X | X | X | X | X | 3.75 | 0-750 | - | C* | 10.6075/J03F4MZC |
|  | NH0911 | CalCOFI | 2009 | 11/06-11/23 (18) | X | X | X | X | X | 3.75 | 0-750 | - | C* | 10.25921/kb75-zk41 |
|  | NH1001 | CalCOFI | 2010 | 01/12-02/06 (26) | X | X | X | X | X | 3.75 | 0-750 | - | C | 10.25921/7fb3-sx13 |
|  | NH1008 | CalCOFI | 2010 | 07/30-08/17 (19) | X | X | X | X | X | 3.75 | 0-750 | - | C | 10.25921/skfd-cy84 |
|  | NH1010(1011) | CalCOFI | 2010 | 10/28-11/13 (17) | X | X | X | X | X | 3.75 | 0-750 | - | C | 10.25921/1kc8-6090 |
|  | NH1101 | CalCOFI | 2011 | 01/12-02/06 (26) | X | X | X | X | X | 3.75 | 0-1000 | - | C | 10.25921/fd9n-h062 |
|  | NH1107(1108) | CalCOFI | 2011 | 07/27-08/13 (18) | X | X | X | X | X | 3.75 | 0-1000 | - | C* | 10.25921/hkk6-gr50 |
|  | NH1110 | CalCOFI | 2011 | 10/16-11/02 (18) | X | X | X | X | X | 3.75 | 0-1000 | - | C | 10.25921/g3h9-70922 |
|  | NH1201 | CalCOFI | 2012 | 01/27-02/13 (18) | X | X | X | X | X | 3.75 | 0-750 | - | C | 10.25921/sp5c-tf02 |
|  | NH1210 | CalCOFI | 2012 | 10/19-11/05 (18) | X | X | X | X | X | 3.75 | 0-1000 | - | C | 10.25921/zsm2-ds43 |
|  | NH1307 | CalCOFI | 2013 | 07/05-07/22 (18) | X | X | X | X | X | 3.75 | 0-1000 | - | C* | 10.25921/0wp5-y584 |
|  | NH1311 | CalCOFI | 2013 | 11/07-11/25 (19) | X | X | X | X | X | 3.75 | 0-1000 | - | C* | 10.25921/m8h1-mv90 |
|  | NH1407 | CalCOFI | 2014 | 07/06-07/22 (17) | X | X | X | X | X | 3.75 | 0-1000 | - | C | 10.25921/3qac-b540 |
|  | NH1411 | CalCOFI | 2014 | 11/05-11/22 (15) | X | X | X | X | X | 3.75 | 0-1000 | - | C* | 10.6075/J0MC8XCZ |
|  | NH1501 | CalCOFI | 2015 | 01/15-02/07 (24) | X | X | X | X | X | 3.75 | 0-1000 | - | C | 10.6075/J0GQ6W3D |
|  | NH1503 | CalCOFI | 2015 | 04/04-04/20 (18) | X | X | X | X | X | 3.75 | 0-1000 | - | C* | 10.25921/kmqe-4s16 |
| Bell M. Shimada | SH1103 | HAKE | 2011 | 06/26-08/31 (55) | X | X | X | X | X | 9.15 | 0-750 | X | C | 10.7289/V53J39XH |
|  | SH1206 | CCE/SAKE | 2012 | 06/26-08/24 (50) | X | X | X | X | X | 9.15 | 0-750 | X | C | 10.25921/8ncq-6n28 |
|  | SH1305 | SAKE | 2013 | 06/06-08/29 (70) | X | X | X | X | X | 9.15 | 0-750 | X | C | 10.7289/V57942N7 |
|  | SH1401 | CalCOFI | 2014 | 01/29-02/07 (10) | X | X | X | X | X | 9.15 | 0-750 | X | C* | 10.25921/0j1x-tb70 |
|  | SH1404 | CPS | 2014 | 04/16-05/06 (24) | X | X | X | X | X | 9.15 | 0-750 | X | C* | 10.7289/V5JQ0Z6K |
|  | SH1405 | CPS | 2014 | 08/28-09/14 (18) | X | X | X | X | X | 9.15 | 0-750 | X | C* | 10.7289/V50Z718D |
|  | SH1507 | HAKE/SAKE | 2015 | 06/20-09/10 (65) | X | X | X | X | X | 9.15 | 0-750 | - | C | 10.7289/V5D50JZQ |
|  | SH1601 | - | 2016 | 01/08-01/09 (29) | X | X | - | X | - | 9.15 | 0-2000 | X | C* | 10.7289/V5FF3QQV |
|  | SH1604 | CalCOFI | 2016 | 04/01-04/23 (15+7) | X | X | X | X | X | 9.15 | 0-750 | X | C | 10.25921/dd0e-m992 |
|  | SH1610 | - | 2016 | 06/30-08/14 (37) | X | X | X | X | X | 9.15 | 0-700 | X | C* | 10.7289/V55X2786 |
|  | SH1612 | CCE | 2016 | 10/05-10/12 (8) | X | X | X | X | X | 9.15 | 0-2000 | X | C | 10.25921/tnv2-k059 |
| Reuben Lasker | RL1601 | CalCOFI | 2016 | 01/07-01/30 (24) | X | X | X | X | X | 7. | 0-700 | X | C | 10.25921/6dbv-6x16 |
| Melville | CCE0810 | CCE | 2008 | 09/30-10/29 (29) | - | X | X | X | X | 3.4 | 0-750 | - | C | 10.6075/J0765CP3 |
|  | CCE1106 | CCE | 2011 | 06/17-07/16 (22) | - | X | X | X | X | 3.4 | 0-1000 | - | C | 10.6075/J0ZP44F1 |
| MacArthur II | M20907 | CalCOFI | 2009 | 07/13-08/03 (20) | - | X | - | X | X | 4. | 0-500 | - | D | 10.25921/g7dz-bk67 |

## 2 Materials and Methods

### 2.1 Acoustic observations in the CCS

We analyze acoustic observations from Simrad EK60 scientific echo-sounders archived by NOAA’s National Centers for Environmental Information (NCEI) Water Column Sonar Data Archive [52]. We focus on 41 cruises from 2006 through 2016 that sampled the water column to at least 500 m depth, across 8 different ships, for a total of 844 days of observations (see details Table 1). These cruises include stock assessment surveys—such as the integrated acoustic survey of Pacific Hake (HAKE), the Pacific Hake and Pacific Sardine survey (SAKE) [50, 54], and the Coastal Pelagic Species survey (CPS) [49]—as well as long-running programs such as the California Current Ecosystem Long-Term Ecological Research (CCE-LTER) and the California Cooperative Oceanic Fisheries Investigations (CalCOFI) [48].

Echosounder data include up to 5 frequencies from 18 to 200kHz, with differences across programs and cruises (Table 1). We focus on 38 and 120kHz observations, which were sampled by all cruises. Raw echograms were converted to time-dependent vertical mean volume backscattering strength, *S*_*v*_ (*dB re* 1 *m*^−1^) at various spatial and temporal resolutions using the MATLAB Echolab MATLAB toolkit developed by Rick Towler. Updated calibration parameters were applied to convert raw power to *S*_*v*_ when available (see calibration status in Table 1).

### 2.2 Data processing

Raw echograms include artifacts and non-biological signals such as impulsive noise, amplification and attenuation noise [55], background noise [56], and echoes from the seafloor. We remove these signals by applying the suite of filters described in [53] (see “Filters” in Fig. 1), correcting *S*_*v*_ for sound absorption and, when possible, transducer motion, and bin the processed echograms onto a regular grid of 1 m depth and 30 s time intervals for subsequent analysis. The same set of parameters are applied to all cruises (see Supplementary Materials S1). We evaluate the suitability of this approach by comparing our results with publicly available processed echograms (see Supplementary Materials S2).

**Figure 1:**
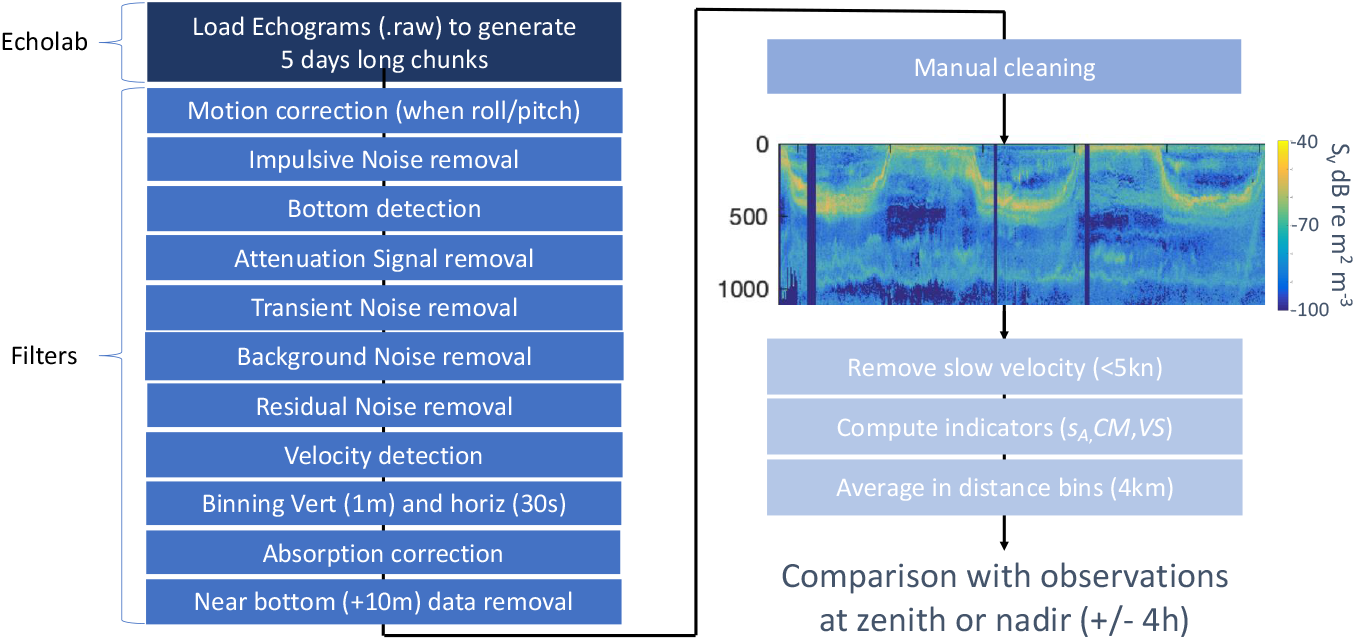
Schematic of the processing steps from raw EK60 echograms to MTL indicators.

Several echograms are affected by additional non-biological signals that are not corrected by the previous filters. These include false bottoms [52], echoes from hydrographic sampling devices such as CTD rosettes, and noise due to interference from shipboard electronic instruments. We inspect all cruises manually and remove intervals affected by these acoustic artifacts (“Manual cleaning” in Fig. 1).

### 2.3 Indicators of MTL distribution

We focus on echogram sections collected while the ship is underway, discarding data acquired at cruising speeds below 5 kn, following standard practice in acoustic-trawl assessments [57, 38]. We then determine a suite of indicators of MTL distribution based on processed *S*_*v*_ profiles (summarized in Table 2) averaged over constant cruising-distance bins (see Indicators in Fig. 1).

**Table 2:**
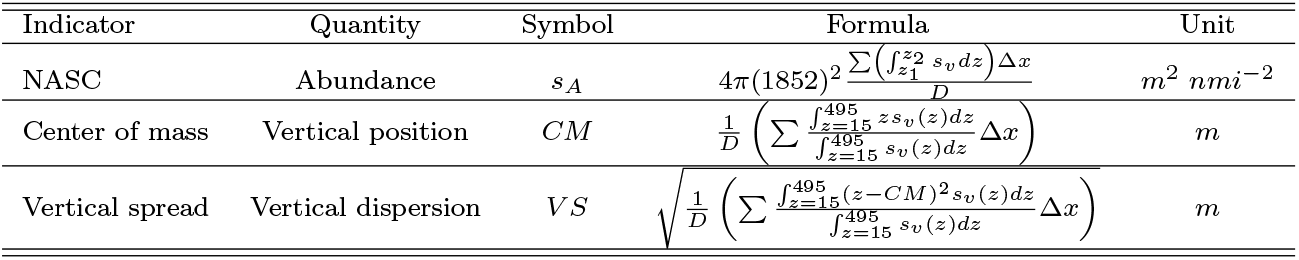
Summary of acoustic indicators of community-level MTL vertical distribution computed from gridded volume backscattering coefficients 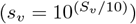 averaged over *D* = ∑Δ*x* = 4 km distance intervals.

First, we compute the Nautical Area Scattering Coefficients (NASC), or *s*_*A*_ (*m*^2^ *nmi*^−2^) [58], over depth strata spanning 15-495 m, defined as:

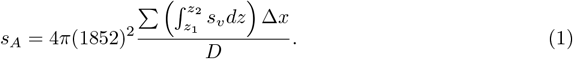

Here, 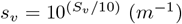 is the volume backscattering coefficient; following vertical integration, values are averaged over distance intervals of width *D* = ∑Δ*x* = 4 km, where Δ*x* is the distance traveled during successive 30 s sampling periods. Depth ranges [*z*_1_ − *z*_2_] are defined as [15 − 175], [175 − 335], [335 − 495] m to coarsely differentiate vertical communities, and as [15 − 110], [110 − 205], [205 − 300], [300 − 395], [395 − 490] m for finer-scale comparisons. To minimize the influence of diel vertical migrations, we evaluate *s*_*A*_ around zenith (± 4*h*) or nadir (± 4*h*), discarding all observations outside these time windows and analyzing daytime and nighttime patterns separately to isolate distinct fish communities. NASC (*s*_*A*_) observations are averaged monthly on a 1/4° grid for analysis.

Computations are based on *S*_*v*_ profiles at 38 kHz and 120 kHz, which primarily sample fish [52]. We use multi-frequency signals to coarsely distinguish dominant organisms types, namely fish with swimbladder and zooplankton [59]. In the upper ocean, where both 38 kHz and 120 kHz data are available, we compute the difference in volume backscattering strength, Δ*S*_*v* 120−38 *kHz*_ = *S*_*v* 120 *kHz*_ − *S*_*v* 38 *kHz*_, for each binned echogram cell. Cells with −9.0 ≤ Δ*S*_*v* 120−38 *kHz*_ < 0. are classified as ‘fish’, and those with 0 ≤ Δ*S*_*v* 120−38 *kHz*_ < 16.5 as ‘zooplankton’. All other signals (Δ*S*_*v* 120−38 *kHz*_ < −9.0 and Δ*S*_*v* 120−38 *kHz*_ > 16.5) are classified as ‘other’ and excluded from further analysis. This classification, derived by comparing Δ*S*_*v* 120−38 *kHz*_ with multi-frequency indices (MFI) computed from all five frequencies, provides a good approximation of MFI-based organism types when only 120 and 38 kHz data are available (see Supplementary Materials S3). In deeper layers (> 175 m), where measurements are limited to 38 kHz, we apply a single “mesopelagic fish” category.

Observations collected during daytime, when vertically migrating organisms reside at depth, allow separation of epipelagic and mesopelagic communities. Differences between day- and night-time observations at depth allow to further partition mesopelagic backscatter (Fig. 3) into migrant (*s*_*A,MM*_) and resident (*s*_*A,MR*_) communities. Accordingly, we define the following acoustic functional groups:

- Zooplankton (*s*_*A,Z*_): *s*_*A*_ from 120 kHz daytime echograms where 0 ≤ Δ*S*_*v* 120−38 *kHz*_ < 16.5 within depth layer [15 − 175] m.
- Epipelagic fish (*s*_*A,F*_): *s*_*A*_ from 38 kHz daytime echograms where −9 ≤ Δ*S*_*v* 120−38 *kHz*_ < 0 within depth layer [15 − 175] m.
- Mesopelagic fish (*s*_*A,M*_): *s*_*A*_ from 38 kHz daytime echograms within depth layers [175 − 335] and [335 − 495] m.

We further separate migrant and resident mesopelagic fish as:

- Migrant mesopelagic fish (*s*_*A,MM*_): nighttime-daytime difference in *s*_*A*_ from 38 kHz echograms where −9 ≤ Δ*S*_*v* 120−38 *kHz*_ < 0 within depth layer [15 − 175] m.
- Resident mesopelagic fish (*s*_*A,MR*_): *s*_*A*_ in 38 kHz nighttime echograms within depth layers [175 − 335] and [335 − 495] m.

Hereafter, we discuss the variability of these MTL groups, recognizing that this approach only allows a coarse, acoustically derived characterization of community composition. We also compute total *s*_*A*_ in 38 kHz echograms over depth strata between 15-495 m to examine how variability propagates from epipelagic to mesopelagic layers.

Along these coarse MTL abundance proxies, we calculate two additional indicators of vertical distribution. The center of mass *CM* (m) provides a measure of the backscatter-weighted mean depth of organisms in the water column. We follow the definition by [60], taking averages over *D* = ∑Δ*x* = 4 km distance intervals:

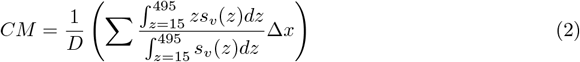

where the volume backscattering coefficients *s*_*v*_(*z*) consist of gridded observations at 1 m vertical resolution.

The vertical spread *V S* (m), also averaged over 4 km distance intervals, quantifies the dispersion of MTLs relative to the center of mass:

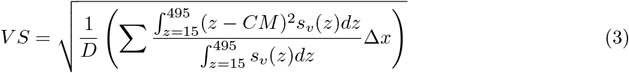

We compute these indicators over the full depth range *z* ∈ [15, 495] m using 38 kHz data, and we average them monthly on a 1/4° grid for subsequent analysis.

### 2.4 Environmental predictors

To characterize environmental drivers of MTL heterogeneity, we compare acoustic indicators (Table 2) to 14 co-located physical and biogeochemical variables from remote sensing [61, 62], ocean reanalysis [63], and interior reconstructions [64] (see Table 3). Environmental variables are linearly interpolated to the mean latitude, longitude, and time of acoustic observations binned at 4 km intervals.

**Table 3:**
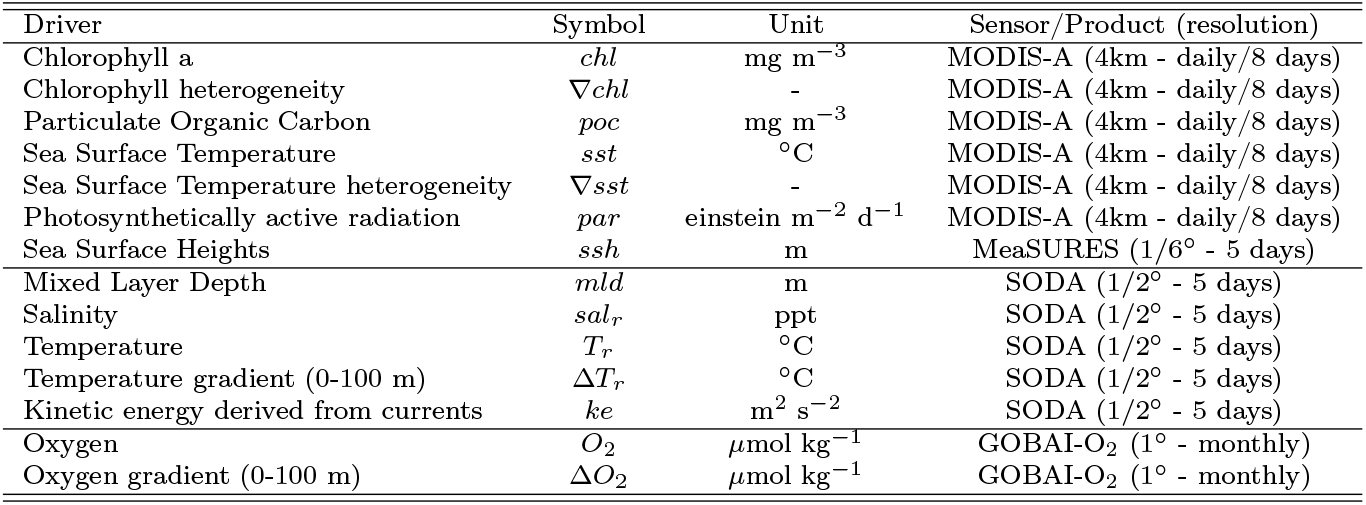
Environmental observation collocated with indicators for MTLs derived from acoustic observation.

First, we use satellite retrievals of surface chlorophyll (*chl* in mg m^−3^), particulate organic carbon (*poc* in mg m^−3^), sea surface temperature (*sst*, in °C) and photosynthetically active radiation (*par* in einstein m^−2^ d^−1^) from MODIS-A [61]. To assess the influence of fronts and filaments, we apply Sobel operators to *sst* and *chl* to compute horizontal gradients (∇*sst* and ∇*chl*), used as proxies for surface habitat heterogeneity associated with oceanic fronts [65]. We use these two indices separately, as temperature and chlorophyll fronts may differentially affect biological processes and do not necessarily co-occur [66]. MODIS-A fields are used at daily resolution when available, and at 8-day resolution otherwise. Where cloud cover produces gaps in daily observations, we derive daily estimates by linearly interpolating between the two nearest 8-day composites. We additionally co-locate sea surface height anomalies (*ssh* in *m*) from the MeaSURES product at 1/6° and 5 days resolution [62].

Second, to account for the effects of currents and water column structure, we use the three-dimensional data-assimilative SODA reanalysis [63] at 1/2° resolution and 5 days frequency, including mixed layer depth (*mld*, in m), sea surface temperature (*T*_*r*_ in °C), and sea surface salinity (*sal*_*r*_ in ppt). We also extract the vertical temperature gradient (Δ*T* in °C between the surface and 100 m depth) and the kinetic energy of the geostrophic current (*ke* = (*u*^2^ + *v*^2^)/2 in m^2^ s^−2^, with *u* and *v* the zonal and meridional components), averaged over the upper 100 m.

Finally, to assess the effect of oxygen, which is known to affect MTL distributions, particularly in the mesopelagic [37, 39], we include both surface oxygen (O_2_ in *µ*mol kg^−1^) and the vertical oxygen gradient (Δ O_2_ in *µ*mol kg^−1^ in the upper 100 m) based on the monthly GOBAI-O_2_ product [64].

For the analysis, the 14 environmental variables (Table 3), aggregated in 4 km bins, are averaged monthly and onto a 1/4° grid spanning the CCS, consistent with the acoustic observations.

### 2.5 Analysis of MTL variability

We process separate subsets of the data to assess the variability of MTLs along three major spatiotemporal gradients. At sub-regional scale, we characterize (1) cross-shore distribution and (2) seasonality of MTLs in Southern California, south of 35°N—an area with extensive offshore and seasonal sampling [48]. At coast-wide scale, we quantify (3) meridional variations along a coastal band extending to 370 km offshore, tracking acoustic observations near the period of peak upwelling, consistent with the northward progression of summertime surveys [50]. To reduce biases arising from uneven seasonal sampling, we retain observations collected from one month before to two months after the peak month of the Biologically Effective Upwelling Transport Index (BEUTI) [32] aggregated into 1° latitude bins.

Along each gradient, for each MTL group, we report the mean values and range of variability (25th and 75th percentiles) of the gridded acoustic MTL indicators. We also quantify correlations between these gridded indicators and collocated environmental features by fitting linear models for each indicator-feature pair, separately controlling for cross-shore distance, month, or latitude as a fixed effect. Data are standardized using z-score normalization, and we report the regression coefficient and p-values as estimates of the strength and significance of each correlation.

## 3 Results

### 3.1 Validation of the acoustic processing approach

To evaluate the robustness of our processing, we compare our echograms with publicly available datasets [67]. For 14 of the 41 transects (see Supplementary Materials S2), our processing reproduces previously published observations with high fidelity in the upper ocean, with most transects showing *R*^2^ > 0.95 and RMSE < 1.5 dB. Larger discrepancies emerge at depth, likely reflecting differences in background noise removal. For a subset of cruises (e.g., NH1001, MF1004, NH1011, NH1210), our approach also reproduces deep backscatter with good agreement, supporting the overall soundness of the processing.

### 3.2 Spatiotemporal data coverage

After processing the 41 acoustic transects, applying spatial averaging, and removing twilight periods, we obtain 14, 209 daytime and 11, 331 nighttime observations aggregated within 4-km bins between 2006 and 2016 (Fig. 2a). When subsequently averaged monthly on a 1/4° grid, these yield 2, 802 daytime and 2, 407 nighttime independent data points.

**Figure 2:**
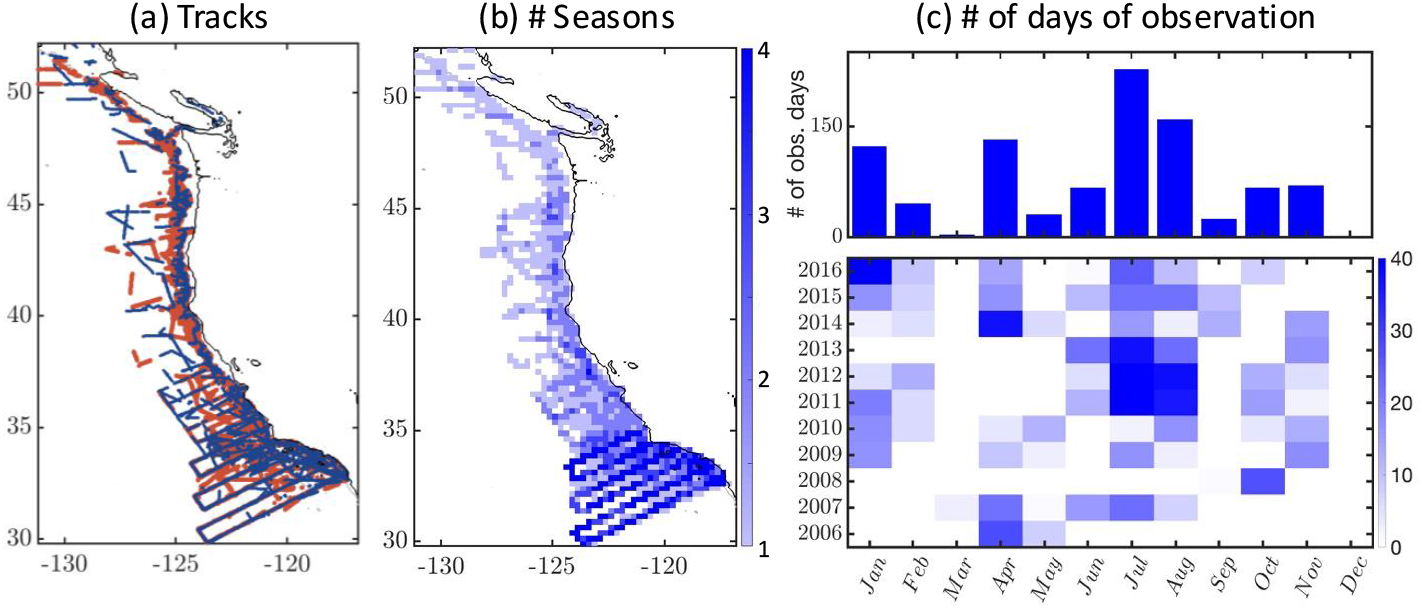
Spatiotemporal data coverage in the California Current System from 2006 to 2016: (a) Day (red) and night (blue) observations along cruise tracks; (b) Number of seasons represented, with data aggregated on a 1/4° grid; (c) Monthly number of observation days, shown as an aggregate histogram for all years (top) and for individual years (bottom).

Coverage is extensive in Southern California, where observations extend hundreds of km offshore along CalCOFI lines, whereas farther north observations are largely confined to the continental shelf and shelf break. Southern California is also sampled consistently year-round, with all four seasons represented, primarily through CalCOFI surveys (Fig. 2b). In the Central and Northern CCS, at most a single season is sampled consistently by the HAKE and SAKE surveys [50].

Interannually, summer months (June to August) are sampled across most years, while winter, spring, and fall observations occur more intermittently, with fewer gaps after 2009 (Fig. 2c). Given the importance of interannual variability in the CCS, the irregular distribution of acoustic observations motivates our focus on subsets of the dataset that best resolve MTL distributions and linkages among communities.

### 3.3 Epipelagic to mesopelagic backscatter distributions (*s*_*A,Z/F/M*_)

#### 3.3.1 Cross-shore variability in Southern California

Analysis of aggregated sonar data reveals consistent spatiotemporal patterns across acoustic functional groups. Daytime observations show a rapid decline in both zooplankton and epipelagic fish backscatter with distance from shore, whereas mesopelagic fish remain more homogeneously distributed along the cross-shore direction (Fig. 3a). The decline occurs over a shorter spatial scale for zooplankton (∼ 100km) than for epipelagic fish (∼ 200km). The range of variability (shown by the envelopes in Fig. 3a), arising from both seasonal and interannual fluctuations, is larger for fish than for zooplankton and decreases rapidly offshore for epipelagic organisms, but not for mesopelagic fish (Fig. 3a). Note that these patterns are conserved when we only consider acoustic transects with complete calibration information (calibration category C in Tab. 1; see Supplementary Materials S4)

**Figure 3:**
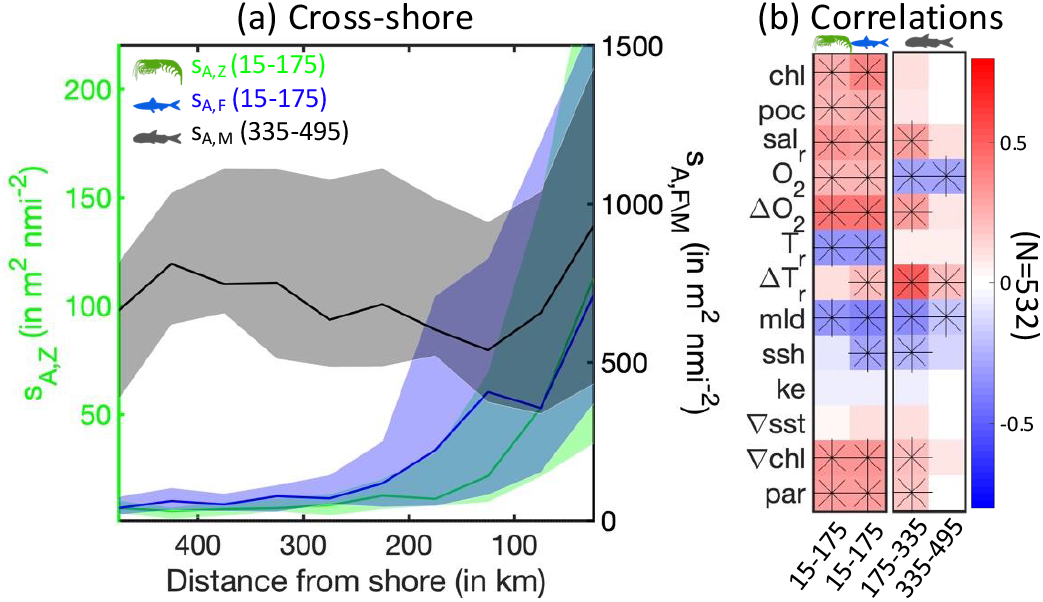
Cross-shore backscatter variability from epipelagic to mesopelagic layers in Southern California. (a) Daytime nautical area scattering coefficient (NASC, or *s*_*A*_, in *m*^2^ *nmi*^−2^) as a function of distance to shore (in *km*) for acoustically defined epipelagic (15-175 m) zooplankton (*s*_*A,Z*_, green) and fish (*s*_*A,F*_, blue) and mesopelagic (335-495 m) fish (*s*_*A,M*_, black). Solid lines show the median across the 11 years of observations, and envelopes the 25th-75th percentile range. (b) Correlations between environmental variables and daytime NASC, based on linear models treating cross-shore variability as a fixed effect. Asterisks (∗) denote correlations with p-value < 0.001; *N* indicates the number of data points.

The cross-shore MTL distribution correlates with multiple environmental variables (Fig. 3b), reflecting strong gradients from coastal to open ocean waters characteristic of the CCS (shown for each variable in Supplementary Materials S5). Across communities and depth layers, most correlations retain the same sign (Fig. 3b), suggesting that MTLs co-vary from epipelagic to mesopelagic habitats. Only the deepest mesopelagic layer appears more weakly connected to shallower layers, as suggested by the loss of significance for many correlations and the flattening of the cross-shore distribution at depth (Fig. 3a, black).

The environmental variables that correlate most strongly with acoustic backscatter are those related to vertical habitat (Δ*O*_2_, Δ*T*_*r*_ and *mld*) and water mass (*sal*_*r*_ and *O*_2_). Productivity (*chl* and *poc*) and solar radiation (*par*) also correlate significantly with surface MTL communities. In contrast, variables related to currents and fine scale ocean features, such as kinetic energy (*ke*), sea surface heights (*ssh*), and horizontal gradients (∇*sst*), show the weakest correlations.

#### 3.3.2 Seasonal variability in Southern California

Daytime acoustic observations reveal a consistent seasonal succession following spring upwelling, with acoustic backscatter peaking first for zooplankton, then epipelagic fish, and finally mesopelagic fish (Fig. 4a). This pattern is consistent with lagged biological responses to peak upwelling, as indicated by the maximum in BEUTI. For the mesopelagic layer, the available data do not allow for clear distinction between a lag and anticorrelation with upwelling, although a lagged response appears plausible. Intraseasonal to interannual variability, along with along-shore variations, contribute to significant spread around the mean (envelopes in Fig. 4a); yet, the seasonal cycle remains robust, particularly for epipelagic and mesopelagic fish, and is preserved when considering only calibrated transects (see Supplementary Materials S4).

**Figure 4:**
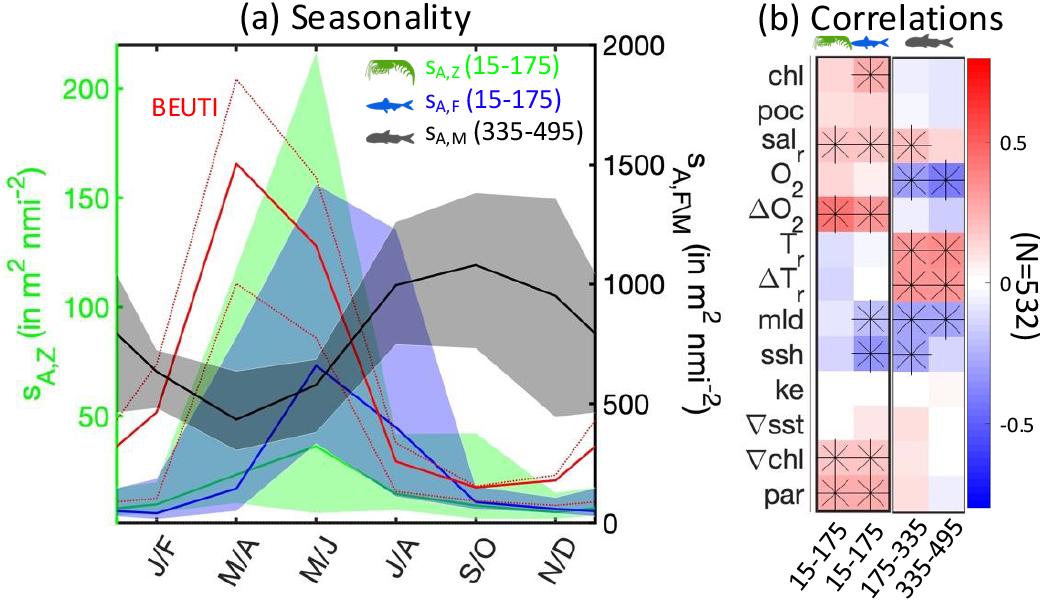
Seasonal backscatter variability from epipelagic to mesopelagic layers in Southern California. (a) Daytime nautical area scattering coefficient (NASC, or *s*_*A*_, in *m*^2^ *nmi*^−2^) as a function of month of the year for surface epipelagic zooplankton *s*_*A,Z*_ (green), fish *s*_*A,F*_ (blue) and mesopelagic fish (335-495 m) *s*_*A,M*_ (black). Solid lines show the median across 11 years of observations, and envelopes the 25th-75th percentiles. The solid red line shows the mean BEUTI index, and the dotted lines indicate the 25th-75th percentiles across years in Southern California [32]. (b) Correlation between environmental variables and daytime NASC, based on linear models where seasonal variability is treated as a fixed effect. Asterisks (∗) denote correlations with p-value < 0.001; *N* indicates the number of data points.

Consistent with the phase shift in the seasonal cycle across MTL communities, correlations with environmental properties vary in both significance and sign across groups and depths (Fig. 4b; seasonal cycles for environmental variables are shown in Supplementary Material S6). Epipelagic fish correlate most strongly with features that vary early in the upwelling season (Δ*O*_2_, *ssh*, and *par*), whereas mesopelagic fish show stronger correlations with subsurface water-mass properties (*sal*_*r*_, *O*_2_, and *T*_*r*_) and post-upwelling stratification (Δ*T*_*r*_ and *mld*) (Figure 4b). Zooplankton exhibits weaker correlations overall, with the stronger relationships found for variables related to productivity (*chl, poc*, and *par*) and vertical habitat compression (Δ*O*_2_). Similar to the cross-shore analysis, seasonal correlations are weakest for features related to currents and fine-scale structures (*ke*, ∇*sst*, and ∇*chl*).

#### 3.3.3 MTL succession

Together, cross-shore and seasonal backscatter variability indicate consistent shifts in the distribution of MTLs from the epipelagic to the mesopelagic zones. Surface NASC (*s*_*A*_) anomalies for zooplankton and 38 kHz signals (Fig. 5), relative to the depth layer mean, show a progressive deepening of backscatter along the cross-shore direction, from the shallow coastal epipelagic maximum (see *s*_*A*_ for the 15 − 110 m layer Fig. 5a) to the deep mesopelagic maximum (395 – 490 m) approximately 500 km offshore (Fig. 5a). On seasonal timescales, the spring epipelagic maximum progressively shifts to deeper layers, peaking between fall and winter in the mesopelagic zone (Fig. 5b). When vertically migrating MTL swim to the surface at night, variability in the mesopelagic layers is attenuated (compare top and bottom panels in Fig. 5).

**Figure 5:**
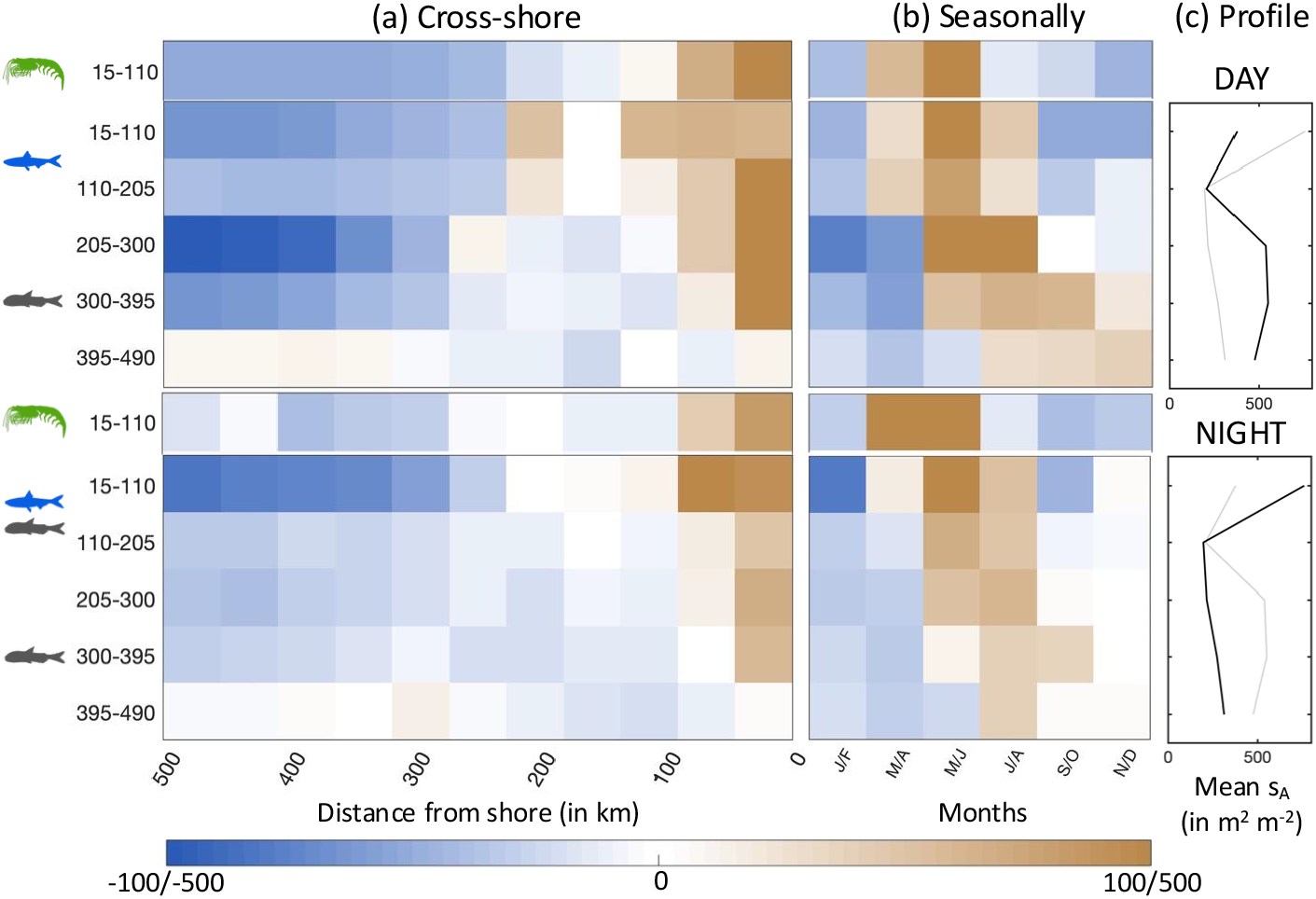
Backscatter variability across vertical layers in Southern California. (a) Day and night-time cross-shore nautical area scattering coefficient (NASC, or *s*_*A*_, in *m*^2^ *nmi*^−2^) anomalies for surface zooplankton (120 kHz) and water-column fish (38 kHz), calculated for 95 m depth intervals and 50 km distance bins, relative to the mean NASC within each layer. (b) Day and nighttime seasonal *s*_*A*_ NASC anomalies for surface zooplankton (120 kHz) and water-column fish (38 kHz), calculated for 95 m depth intervals and 2-month temporal bins, relative to the mean NASC within each layer. (c) Mean daytime and nighttime NASC profiles by depth interval at 38 kHz. In panels (a) and (b), the color range is [−100, 100] for zooplankton and [−500, 500] for fish.

#### 3.3.4 Alongshore variability

Across latitudes, acoustic observations during peak upwelling reveal shifts in distribution between zooplankton and fish, as well as between epipelagic and mesopelagic layers. For epipelagic communities, observations show two backscatter maxima around 35°N and 43°N (see Fig. 6a, showing epipelagic fish), with the southern zooplankton peak slightly displaced northward relative to fish (compare green and blue lines in Fig. 6b; see also Supplementary Materials S4 for calibrated transects only). Large envelopes around the mean reflect significant interannual variability. The zooplankton distribution is more irregular but consistent with localized hotspots associated with convergence and retention zones [27]. The northern maximum is particularly intense in mesopelagic layers and displaced northward to approximately 45°N (compare blue and black lines in Fig. 6b). Notably, these bimodal distributions are offset relative to the single peak in nutrient upwelling centered around 39°N, as indicated by BEUTI (Fig. 6a).

**Figure 6:**
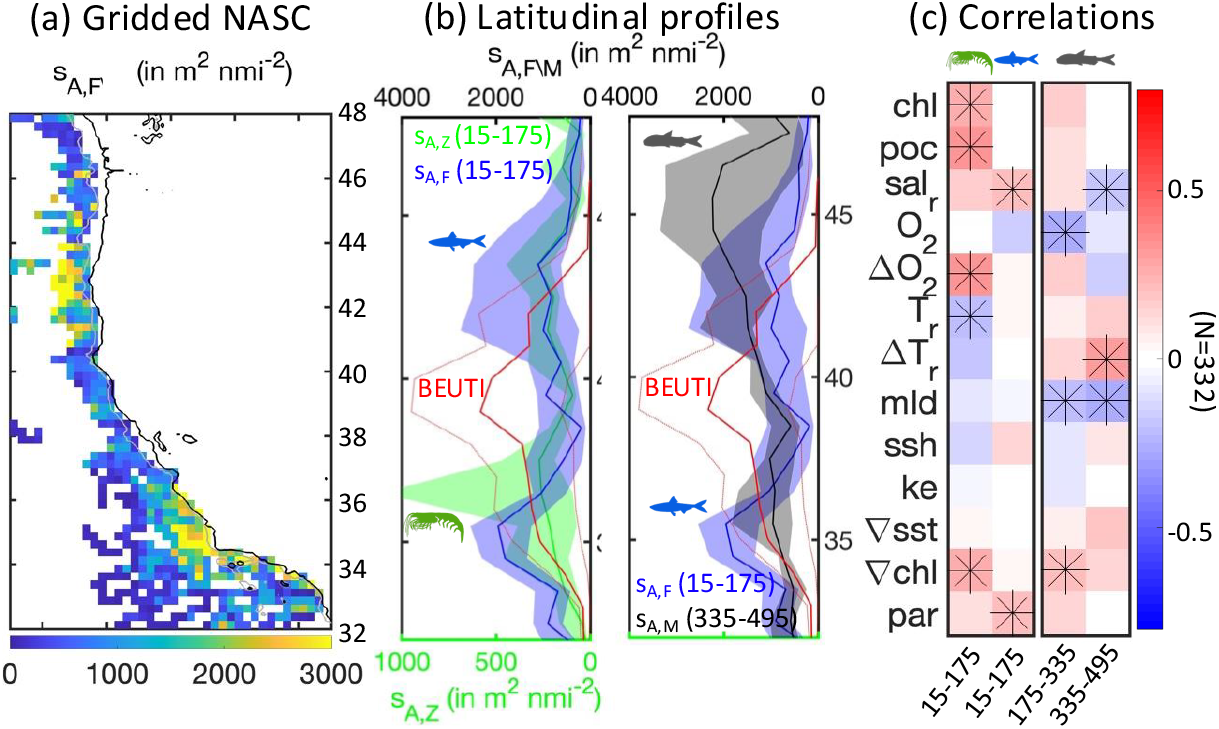
Along-shore backscatter variability from epipelagic to mesopelagic layers. (a) Daytime nautical area scattering coefficient (NASC, or *s*_*A*_, in *m*^2^ *nmi*^−2^) as a function of latitude (in °), mapped for epipelagic fish. (b) Profiles of daytime NASC for epipelagic zooplankton *s*_*A,Z*_ (green), epipelagic fish *s*_*A,F*_ (blue), and mesopelagic (335-495m) fish, *s*_*A,M*_ (black). In the profiles, solid lines show the 50th percentile across the 11 years of observations, and envelopes the 25th-75th percentiles. The solid red line shows the mean BEUTI across years and latitudes; dotted lines the 25th-75th percentiles across years [32]. (c) Correlation between environmental variables and daytime NASC, based on linear models where alongshore variability is treated as a fixed effect. Asterisks (∗) denote correlations with p-value < 0.001; *N* indicates the number of data points.

Alongshore, acoustic backscatter shows similar but weaker correlations with environmental predictors (Fig. 6c) than those observed for cross-shore and seasonal variability in the southern domain (meridional distributions of environmental variables are shown in Supplementary Materials S7). The broad latitudinal extent of the analysis spans multiple water masses and species assemblages within the California Current, likely complicating the identification of clear relationships with environmental drivers. Epipelagic zooplankton show stronger correlations with productivity-related variables (*chl* and *poc*) and, to a lesser extent, with vertical O_2_ gradients (Δ*O*_2_), suggesting a response to upwelling intensity and habitat compression. In contrast, mesopelagic fish correlate more strongly with water-mass-related variables (Δ*T*_*r*_, *O*_2_, *sal*_*r*_ and *mld*).

### 3.4 Mesopelagic migrant and resident backscatter (*s*_*A,MM/MR*_)

Separation of mesopelagic fish into migrant and resident communities shows that, whereas the former declines gradually offshore, the latter remains more uniform (Fig. 7a). The spread is larger for migrant MTLs, potentially reflecting larger intraseasonal to interannual variability, similar to epipelagic groups. Seasonal variability of migrant and resident communities is comparable to that of the daytime mesopelagic backscatter (Fig. 4), suggesting a synchronous response, with both exhibiting the same lagged seasonal cycle relative to upwelling and surface MTL (Fig. 7b). The sparseness of co-located day–night observations along the full coast—required to estimate backscatter from migrant mesopelagic fish—precludes identification of clear alongshore patterns across migrant and resident mesopelagic communities.

**Figure 7:**
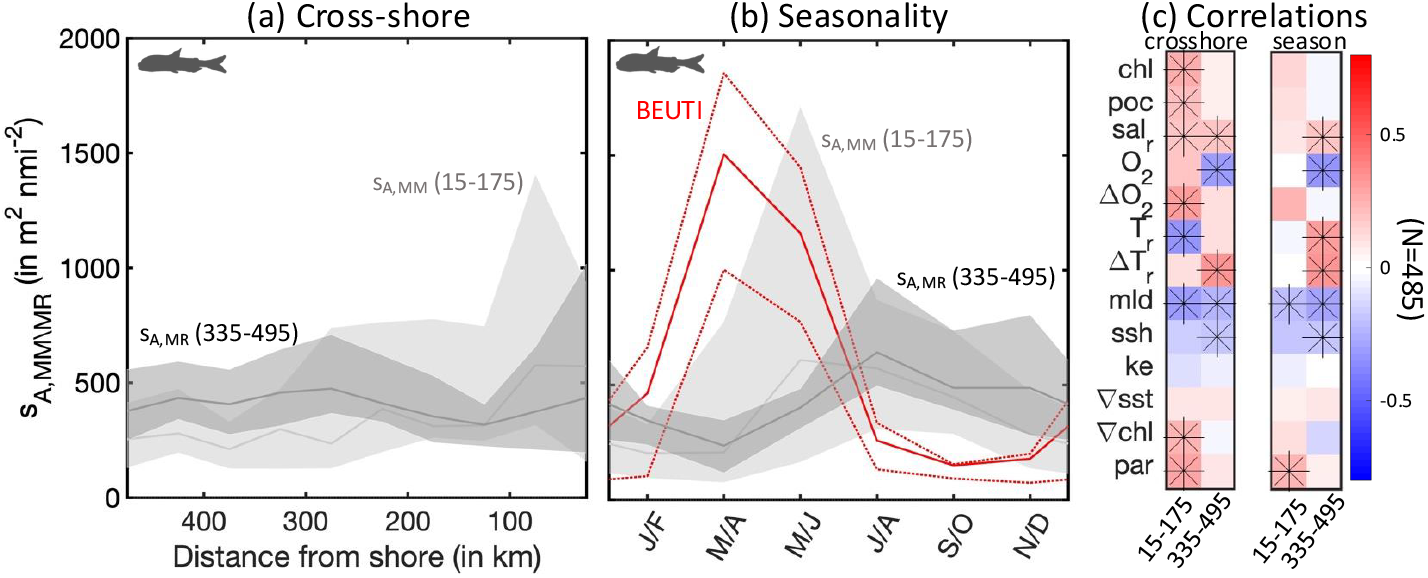
Variability of migrating and resident mesopelagic MTLs. (a) Cross-shore variability of the nautical area scattering coefficient (NASC, in *m*^2^ *nmi*^−2^) for migrating mesopelagic fish (*s*_*A,MM*_, light gray) and resident mesopelagic fish (*s*_*A,MR*_, dark gray). (b) Seasonal variability in NASC for migrating mesopelagic fish (*s*_*A,MM*_) and resident mesopelagic fish (*s*_*A,MR*_). In panels (a,b), solid lines show the 50th percentile across the 11 years of observations, and envelopes the 25th-75th percentiles. The solid red line in panel (b) shows BEUTI averaged across years and latitudes; dotted lines the 25th-75th percentiles across years [32]. (c) Correlation between environmental variables and mesopelagic NASC, based on linear models in which seasonal or alongshore variability are treated as a fixed effect. Asterisks (∗) denote correlations with p-value < 0.001; *N* indicates the number of data points for the cross-shore and seasonal correlations..

Cross-shore and seasonal correlations with environmental properties suggest differences in the dominant drivers influencing migrant versus resident mesopelagic communities (Fig. 7c). Overall, variability of the migrant community—which spends part of the day in the epipelagic zone— correlates with similar features to those of epipelagic groups (compare Figs. 3b and 4b, with surface 15 − 175 correlations in Fig. 7c). For the resident mesopelagic community (layer 335 − 495 in Fig. 7c), correlations instead highlight a dominant link with vertical habitat structure (Δ*T*_*r*_, *mld* and *ssh*) and water mass properties (*sal*_*r*_ and *O*_2_).

### 3.5 Vertical MTL distribution

Along the cross-shore direction, the center of mass of acoustic backscatter (Table 2) deepens by about 150 m between the coast and the open ocean, indicating a progressive downward shift of MTL communities, whereas the vertical spread remains nearly uniform, only slightly declining offshore (Fig. 8a).

**Figure 8:**
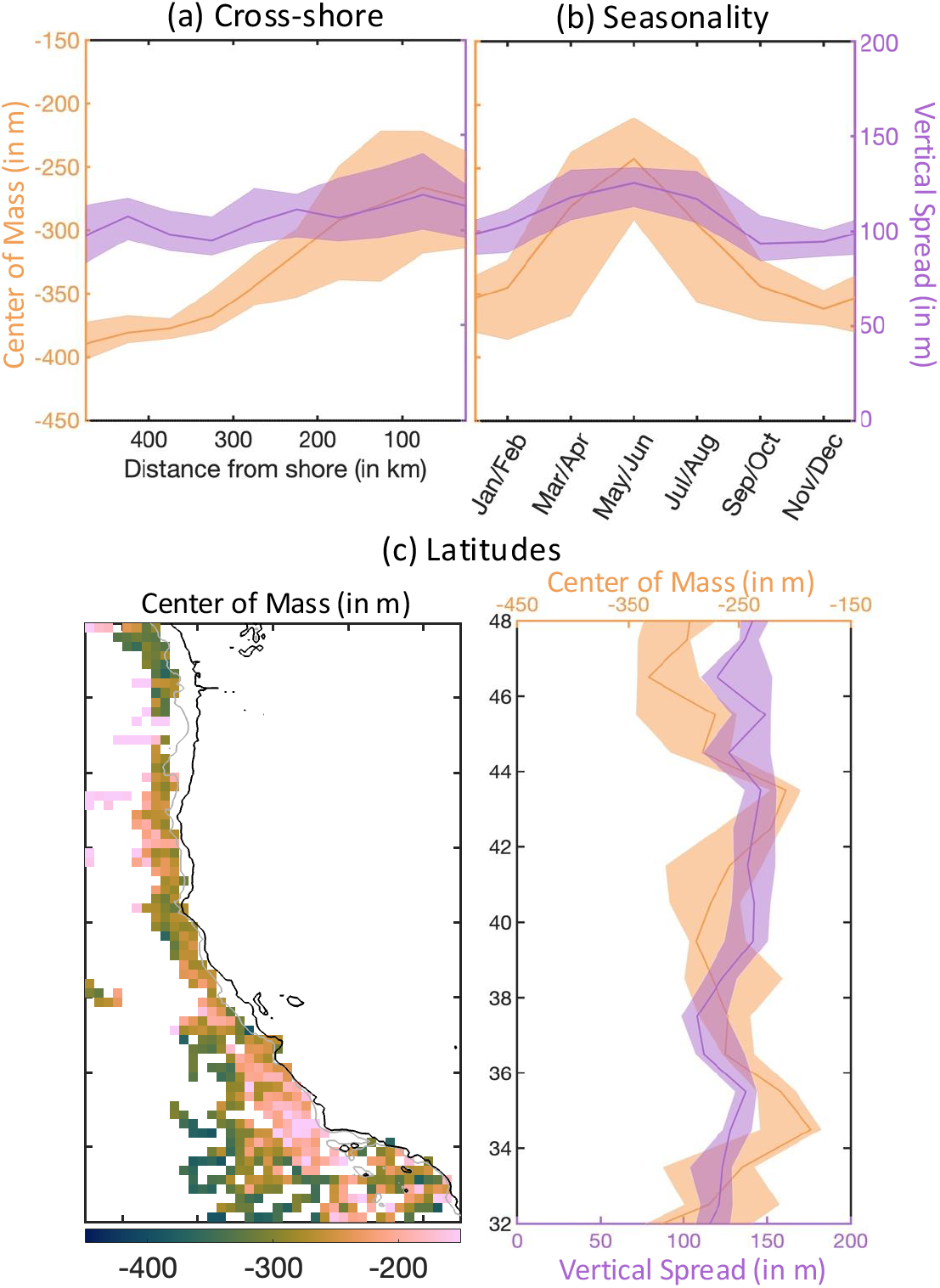
Daytime variability of the center of mass (orange, *m*) and vertical spread (purple, *m*) of acoustic backscatter at 38 kHz, calculated for the upper 500 m. (a) Cross-shore distribution. (b) Seasonal variability. (c) Map of the center of mass and mean along-shore distribution of center of mass and vertical spread. In each profile in panels (a)-(c), the solid lines show medians across the 11 years of observations, and envelopes the 25th-75th percentiles.

Seasonally, the center of mass shoals by approximately 100 m from its deepest position in winter (∼ 350 m) to its shallowest in spring (∼ 250 m, Fig. 8b). This is consistent with increasing epipelagic backscatter during upwelling (see Fig. 4a). The shoaling is accompanied by a slight increase in vertical spread (Fig. 8b and Fig. 4a).

Along the coast, regions of enhanced epipelagic backscatter are characterized by shallower center of mass, with the shallowest depths (less than 200 m) observed just offshore of the shelf break near 34°N and 43°N (Fig. 8c). Overall, the center of mass deepens by ∼ 100 m northward, while changes in vertical spread are less clearly defined.

Indicators of the vertical distribution show the strongest correlations with vertical oxygen gradients (Δ*O*_2_) and light (*par*), whereas other variables show weaker or less consistent correlations once averaged along the three environmental gradients (see Supplementary Materials S8).

## 4 Discussion

Drawing on 11 years of multifrequency fisheries acoustic observations, we evaluate the spatiotemporal distribution of MTL communities in the California Current System. Although a full multi-frequency index [59] cannot be applied because of data gaps, a simplified classification based on backscatter at 38 and 120 kHz, combined with day-night comparisons, enables definition of 4 coarsely resolved functional groups representing epipelagic zooplankton, epipelagic fish, and migrant and resident mesopelagic fish. While backscatter cannot be directly translated into biomass or abundance without accurate knowledge of community composition [68, 69], these simplified proxies provide a synoptic, quantitative characterization of large-scale, community-level patterns and variability in MTL [3, 70, 71, 72]. We discuss caveats and limitations of this approach in a dedicated section below.

Despite heterogeneous sampling and the coarse nature of our acoustic classifications, coherent patterns emerge along three major gradients: cross-shore, seasonal, and alongshore (Figs. 3, 4, 6 and 7). These patterns are broadly consistent with, and extend, a succession-based view of ecosystem responses to seasonal upwelling in the CCS [21, 23, 29]. By investigating relationships with co-located environmental variables, we assess the role of vertical habitat structure and water-mass properties in shaping these distributions. The following sections interpret these results within their ecological context, as summarized in Table 4.

**Table 4:** Ecosystem level pattern of MTL variability and driving mechanisms.

| Gradient | MTL | Pattern of variability | Potential mechanisms |
| --- | --- | --- | --- |
| Cross-shore | Epipelagic | Coastal maximum with strong cross-shore attenuation | M1: Coastal upwelling fuels productivity that propagate towards larger trophic levels and attenuates as distributed offshore by physical transport. |
|  | Mesopelagic | Homogeneous cross-shore distribution | M2: Lower metabolic demands reduce dependence on surface productivity, allowing mesopelagic distributions to remain more homogeneous. |
| Seasonal | Epipelagic | Lagged seasonal cycles from zooplankton to epipelagic fish | M3: Nutrient rich upwelling initiates a trophic succession, with lagged peaks from zooplankton to epipelagic fish.<br>M4: Epipelagic fish migrate in response to habitat compression |
|  | Mesopelagic | Seasonal cycle out of phase with epipelagic groups | M5: Shoaling of hypoxic waters during upwelling displace mesopelagic communities.<br>M6: Penetration of warmer and less oxygenated tropical waters seasonally shifts dominant mesopelagic assemblages. |
| Latitudinal | Epipelagic | Multiple local maxima for zooplankton, two broader maxima for epipelagic fish | M7: Centers of nutrient rich upwelling and retention regions lead to higher MTL abundance.<br>M8: Modulations linked to the width of the continental shelf and pelagic vs. demersal routing of primary production. |
|  | Mesopelagic | Two broad maxima shifted compared to the epipelagic fish | M9: Seasonal shift of dominant communities as upwelling progresses northward during summertime surveys. |

### 4.1 Cross-shore community succession

Epipelagic backscatter decreases from the coast to the open ocean, with progressively more uniform cross-shore distributions at higher trophic levels. This structure, well documented for phytoplankton and zooplankton in the CCS [22, 25, 29], is corroborated here by acoustic proxies for epipelagic zooplankton and fish (Fig. 3). Enhanced nearshore backscatter is consistent with higher catches of forage species near the coast [73], and with prior acoustic observations linking elevated MTL backscatter to high primary production [74]. The observed broader offshore extent of epipelagic fish relative to zooplankton is also consistent with prior evidence that planktivorous fish increasingly dominate seaward of the upwelling front compared to macrozooplankton [15].

A plausible mechanism (mechanism M1 in Table 4) is that coastal upwelling and interactions between currents and the shelf break drive high productivity on the continental shelf, while surface Ekman transport and eddies redistribute nutrients and phytoplankton biomass offshore [24]. Slower growth may delay zooplankton and fish biomass peaks farther offshore [29, 30], with convergence zones and eddies providing additional aggregation centers for mobile organisms [75]. Substantial variability can be attributed to interannual expansion and contraction of favorable shallow habitats [34], as well as to changes in upwelling intensity that may favor different forage species [33]. In addition, meridional migrations, common among epipelagic forage fish such as anchovies, can lead to shifts in regional distributions within and outside the Southern California region, rather than producing a simple offshore expansion [18].

In contrast, mesopelagic communities show a more homogeneous cross-shore distribution, albeit with different spreads for migrant and resident components. Together, this implies a greater relative contribution of mesopelagic organisms towards the open ocean as surface groups decline in abundance—a contrast consistent with diet studies indicating an increasing fraction of mesopelagic prey in the stomachs of marine predators caught offshore [46].

Partial decoupling between epipelagic and mesopelagic communities suggests a weaker association of mesopelagic backscatter with coastal productivity (Fig. 7). A plausible mechanism (M2 in Table 4) is reduced metabolic rates in cold, deep waters, which allow mesopelagic organisms to persist under lower resource availability, thereby dampening the imprint of surface productivity gradients. Habitat-related constraints appear more influential for mesopelagic communities (see correlation with Δ*O*_2_, Δ*T*_*r*_, and *mld* in Fig. 3), particularly shoaling of hypoxic waters towards the coast [37]. Meanwhile, mesopelagic communities associated with distinct water masses [39] may differ in species composition, size structure, and acoustic target strength, such that similar backscatter levels do not necessarily imply uniform biomass along the cross-shore direction.

Contrasting gradients in epipelagic and mesopelagic communities are reflected in the progressive deepening of the center of mass with distance from shore (Fig. 8), driven by the combined effects of declining epipelagic backscatter and deepening of mesopelagic layers offshore. However, because the mesopelagic community likely extends below the depth range sampled here (i.e., deeper than 495 m; Fig. 5), the observed cross-shore homogeneity probably represents a conservative view of variability at depth.

### 4.2 Seasonal community succession

Spring upwelling triggers phytoplankton blooms that fuel zooplankton accumulation and ultimately sustain fish biomass peaks later in the season, consistent with the succession observed in epipelagic acoustic backscatter (M3 in Table 4, see also Fig. 4a). The post-peak decline in epipelagic fish may reflect meridional migrations [18] and zonal redistribution as favorable habitats expand offshore— although the steep cross-shore gradient supports alongshore movement—as well as predation mortality. Notably, epipelagic backscatter correlates strongly not only with productivity proxies but also with indicators of vertical habitat structure, such as Δ*O*_2_ and mixed-layer depth (Fig. 4b), suggesting higher abundance during periods of vertical habitat compression, when feeding success may be enhanced (M4 in Table 4).

In contrast, both migrant and resident mesopelagic communities display an opposite seasonal pattern (Fig. 4a), with minimum backscatter during upwelling and a maximum from late summer to fall (Fig. 7b). This phasing is consistent with observations from upward-looking echosounders in Monterey Bay, which show a seasonal disappearance of the resident mesopelagic scattering layer during upwelling [14].

Shoaling of hypoxic waters during upwelling may displace mesopelagic communities offshore (M5 in Table 4), consistent with the observed anticorrelation with *O*_2_, although not with the vertical oxygen gradient Δ*O*_2_ (Figs. 4b and 7c). Alternatively, seasonal shifts in dominant water masses and associated mesopelagic communities—potentially linked to the inflow of warmer, less oxygenated waters of tropical origin—may contribute to the observed backscatter variability (M6 in Table 4). Such water-mass-driven changes in community composition are observed interannually through shifts in mesopelagic ichthyoplankton assemblages [10].

The opposing seasonal phases lead shifts in the mean position of organisms in the water column, which is shallowest during peak spring upwelling, and deepens progressively through summer into fall and winter (Fig. 5). This pattern likely reflects a combination of upwelling-driven changes in mesopelagic habitat and shoaling of isolumes associated with increased turbidity during phytoplankton blooms.

### 4.3 Alongshore variability

Multiple maxima in meridional backscatter are consistent with a heterogeneous distribution of MTLs—from krill to micronekton to top predators [76]. In particular, zooplankton maxima coincide with krill hotspots near approximately 35°, 37° and 41°N [77] (M7 in Table 4), which have been linked to above-average nutrient supply and phytoplankton accumulation within coastal convergence zones [27]. Epipelagic fish show two broader backscatter maxima that are slightly offset from the zooplankton peaks (Fig. 6a). Clear environmental drivers for this pattern are not readily identified; however, weaker backscatter is observed in regions with broader continental shelves (Fig. 6a,b, compare bathymetry with alongshore profiles). This may partly reflect observational bias arising from the exclusion of backscatter in shallow waters. Alternatively, ecological processes such as enhanced routing of primary production to demersal communities may contribute (M8 in Table 4) [78]. Notably, maxima in epipelagic backscatter flank the region of peak upwelling (see BEUTI; Fig. 6b).

Two alongshore maxima are also evident for mesopelagic fish, albeit shifted relative to epipelagic backscatter, and with significantly greater amplitude at higher latitudes (Fig. 6b), consistent with the northward deepening of the center of mass (Fig. 8c).

Identifying dominant drivers of alongshore variability is challenging, in part because this broad latitudinal gradient spans distinct ecosystems and water masses, each characterized by different MTL assemblages [11]. Furthermore, the timing of acoustic surveys and the focus on upwelling-season observations may also influence the identified patterns. Surveys are typically conducted during summer, beginning in the south in June and progressing northward. As a result, southern latitudes are sampled when epipelagic backscatter is near its seasonal peak (Fig. 4), whereas higher latitudes are sampled later in the season, when epipelagic backscatter may have declined and mesopelagic backscatter may be increasing (M9 in Table 4). However, the extent to which this sampling bias contributes to alongshore variability is unclear.

### 4.4 Caveats

Our analysis builds on echograms collected across multiple surveys in the CCS [49, 48, 52, 54]. Although all transects use Simrad EK60 echosounders, data were acquired on different platforms with varying characteristics and, in some cases, partial calibration information (see Table 1). To minimize methodological discrepancies, we applied a single standardized processing framework, following prior work [53]. While this approach might overlook cruise-specific factors, comparisons with previously published echograms indicate minimal differences (see Supplementary Materials S2).

Acoustic observations are routinely used to quantify MTL biomass; however, this requires thorough calibration and community composition data from co-located trawl samples [68, 57]. Here, we do not attempt such pairing, and our approach does not account for variability in acoustic signals due to variations in species composition, life stage, size structure, and behavior [79, 69]. Therefore, the observed acoustic signals can only be interpreted as approximate, qualitative proxies of MTL abundance.

Instead, we focus on relative variations in acoustic backscatter, classified based on a limited set of traits: depth strata, day-night changes, and contrast between low (38 kHz) and high (120 kHz) frequencies. The consistency of recovered patterns with prior CCS studies [74, 14, 73, 15, 27, 46] suggests that this simplified approach captures meaningful ecological information at the community level. Furthermore, the observed patterns are conserved whether considering only fully calibrated transects or all available transects (see Supplementary Materials S4). Incorporating additional knowledge, such as dominant fish assemblages [10, 11] and regional-scale traits [80], could further improve interpretation of MTL variability.

Finally, our approach is constrained by the spatiotemporal coverage of acoustic surveys (Fig. 2). Sampling at higher latitudes is largely limited to summer, introducing gaps that complicate interpretation. For instance, some latitudinal patterns (Fig. 6) may reflect aliasing of seasonal dynamics onto spatial gradients. The identification of significant relationships between MTL backscatter and co-located environmental variables—particularly in the densely sampled Southern California region—suggests that backscatter distributions are, to some extent, predictable. Future work leveraging statistical and machine-learning approaches to extrapolate MTL dynamics into less well-sampled regions may provide additional insight [81], although preliminary efforts highlight substantial challenges [82].

## 5 Conclusions

Our analysis of acoustic observations characterizes variability in zooplankton, epipelagic fish and mesopelagic fish communities in the CCS, revealing three patterns: (1) a cross-shore structure marked by a strong coastal maximum in surface backscatter, with a nearshore zooplankton peak and a broader fish maximum, contrasting with more uniform mesopelagic backscatter; (2) a seasonal succession in which epipelagic backscatter peaks following the onset of upwelling, whereas mesopelagic backscatter reaches a maximum well after upwelling relaxation in fall and winter; and (3) pronounced alongshore heterogeneity, with multiple latitudinal backscatter maxima for each MTL group, offset northward or southward relative to the upwelling core near 39°N. A progressive deepening of mesopelagic backscatter is evident offshore.

These emerge from years of heterogeneous observations and are thus associated with large uncertainty, driven in part by interannual modulations of upwelling. Yet, several aspects of the observed variability match expectations based on previous studies, including the steep offshore decline of epipelagic backscatter [74], lagged seasonal dynamics in mesopelagic layers [14], and alongshore zooplankton hotspots [76, 77, 27]. Taken together, these results suggest an extension of trophic succession dynamics from zooplankton to MTL fish, visible along both cross-shore and seasonal gradients. This extended framework links physical forcing and nutrient supply to phytoplankton and zooplankton biomass, and onward to MTLs, with temporal delays and spatial shifts reflecting, among other factors, organism-specific growth timescales [30], transport by currents [77, 27, 75], and climate-driven variations in habitat characteristics [33, 2, 37].

By combining more than a decade of observations, we mitigate the inherently sporadic nature of fisheries acoustic sampling and identify ecosystem-wide patterns of MTL variability. MTL abundance and spatial distribution [34, 10] exhibit large variations due to interannual variability in both physical drivers and biological responses [44]. Quantifying the effect of interannual forcings on large-scales acoustic patterns will be a critical step toward developing a coherent, community-level view of MTL dynamics and change in the CCS [50, 38].

## 6 Acknowledgments

We thanks S. Bebin, L. Garrobé and J. Marshall for their contributions during preliminary stages of the analysis and for their assistance with data processing. We gratefully acknowledge all contributors to the surveys that collected the acoustic data. This project received funding from US National Aeronautics and Space Administration (NASA), Grants Number 80NSSC21K0420 and 80NSSC25K7430. An AI-based large language model was used to review selected text for clarity and grammar using the following prompt: “The context is a scientific manuscript on mid-trophic levels in the California Current Ecosystem, based on acoustic observations. Improve the following paragraphs or sentences for flow and grammar, keeping as close as possible to the original.”

## 7 Data statement

The acoustic data, processed and binned along survey transects and depth strata with collocated environmental observations, and the scripts to produce the figures and analyses are available at https://doi.org/10.5281/zenodo.19462565.

