## Supplementary Materials for "Epipelagic to mesopelagic variability of acoustic backscatter in the California Current"

April 7, 2026

### S1 Calibration and processing parameters

Table 1: Processing parameters for all filters

| Parameter | Unit | Value used |
|---|---|---|
| Impulsive noise | | |
| Exclusion threshold | $dB\ re\ 1\ m^2\ m^{-3}$ | $-170$ |
| Detection threshold | $dB\ re\ 1\ m^2\ m^{-3}$ | 6 |
| Vertical size of smoothing window | $m$ | 5 |
| Horizontal size of context window | # | 3 |
| Attenuation signal | | |
| Detection threshold | $dB\ re\ 1\ m^2\ m^{-3}$ | 8 |
| Detection percentile | % | 50 |
| Exclude above depth | $m$ | 300 |
| Exclude below depth | $m$ | 500 |
| Horizontal size of context window | # | 300 |
| Transient noise | | |
| Detection threshold | $dB\ re\ 1\ m^2\ m^{-3}$ | 15 |
| Exclusion threshold | $dB\ re\ 1\ m^2\ m^{-3}$ | $-150$ |
| Detection percentile | % | 15 |
| Exclude above depth | $m$ | 250 |
| Vertical size of smoothing window | $m$ | 20 |
| Vertical size of context window | # | 10 |
| Horizontal size of context window | # | 50 |
| Background noise | | |
| Minimum SNR threshold | $dB\ re\ 1$ | 5 |
| Maximum noise threshold | $dB\ re\ 1\ W$ | $-100$ |
| Vertical size of averaging window | # | 15 |
| Horizontal size of averaging window | # | 10 |
| Residual noise | | |
| Exclusion threshold | $dB\ re\ 1\ m^2\ m^{-3}$ | $-160$ |
| Detection threshold | $dB\ re\ 1\ m^2\ m^{-3}$ | $-50$ |
| Boolean data range | $dB\ re\ 1\ m^2\ m^{-3}$ | $[-998; -20]$ |
| Vertical size of context window | # | 7 |
| Horizontal size of context window | # | 7 |

### S2 Evaluation of our processing approach

| | 15-115m | 115-215m | 215-315m | 315-415m | 415-515m |
|---|---|---|---|---|---|
| NH0901 (https://doi.org/10.6075/J0XW4H5R) – $R^2$ (RMSE) | | | | | |
| 18kHz | 0.83 (2.70dB) | 0.76 (3.54dB) | 0.79 (1.70dB) | 0.87 (1.41dB) | 0.79 (2.50dB) |
| 38kHz | 0.43 (4.22dB) | 0. (6.80dB) | 0. (6.95dB) | 0. (5.87dB) | 0. (6.11dB) |
| 70kHz | 0.87 (2.16dB) | 0.33 (5.11dB) | 0.34 (2.97dB) | | |
| 120kHz | 0.78 (3.09dB) | 0.17 (6.24dB) | | | |
| 200kHz | 0.10 (7.24dB) | | | | |
| NH1001 (https://doi.org/10.6075/J0PC30Q0) – $R^2$ (RMSE) | | | | | |
| 18kHz | 0.93 (1.98dB) | 0.99 (0.79dB) | 0.98 (0.53dB) | 0.96 (0.85dB) | 0.88 (1.85dB) |
| 38kHz | 0.94 (1.44dB) | 0.99 (0.57dB) | 0.96 (0.72dB) | 0.80 (1.26dB) | 0.78 (1.43dB) |
| 70kHz | 0.97 (1.10dB) | 0.97 (0.82dB) | 0.83 (1.58dB) | | |
| 120kHz | 0.96 (1.18dB) | 0.76 (2.71dB) | | | |
| 200kHz | 0.82 (2.73dB) | | | | |
| NH1008 (https://doi.org/10.6075/J0DV1H7H) – $R^2$ (RMSE) | | | | | |
| 18kHz | 0.98 (1.41dB) | 0.99 (1.34dB) | 0.97 (1.53dB) | 0.87 (1.46dB) | 0.83 (3.12dB) |
| 38kHz | 0.98 (0.83dB) | 0.94 (2.26dB) | 0.85 (3.15dB) | 0.25 (4.98dB) | 0. (6.23dB) |
| 70kHz | 0.99 (0.55dB) | 0.64 (3.92dB) | 0.(5.87dB) | | |
| 120kHz | 0.73 (3.33dB) | 0.73 (2.38dB) | | | |
| 200kHz | 0.93 (1.44dB) | | | | |
| NH1101 (https://doi.org/10.6075/J05D8Q6J) – $R^2$ (RMSE) | | | | | |
| 18kHz | 0.95 (1.99dB) | 0.97 (1.42dB) | 0.92 (1.35dB) | 0.86 (1.93dB) | 0.77 (4.28dB) |
| 38kHz | 0.95 (1.24dB) | 0.89 (2.32dB) | 0.65 (3.62dB) | 0. (6.35dB) | 0. (9.56dB) |
| 70kHz | 0.96 (1.04dB) | 0.25 (8.99dB) | 0. (11.6dB) | | |
| 120kHz | 0.96 (1.09dB) | 0.19 (7.91dB) | | | |
| 200kHz | 0.93 (1.36dB) | | | | |
| NH1108 (https://doi.org/10.6075/J09Z937P) – $R^2$ (RMSE) | | | | | |
| 18kHz | 0.97 (1.25dB) | 0.98 (1.15dB) | 0.98 (1.00dB) | 0.91 (1.26dB) | 0.81 (3.20dB) |
| 38kHz | 0.96 (1.09dB) | 0.96 (1.76dB) | 0.89 (2.54dB) | 0.27 (4.12dB) | 0. (6.84dB) |
| 70kHz | 0.97 (1.11dB) | 0.45 (9.02dB) | 0. (12.5dB) | | |
| 120kHz | 0.90 (1.92dB) | 0. (16.0dB) | | | |
| 200kHz | 0.86 (2.16dB) | | | | |
| NH1202 (https://doi.org/10.6075/J0HM56S0) – $R^2$ (RMSE) | | | | | |
| 18kHz | 0.93(1.84dB) | 0.97 (1.11dB) | 0.92 (0.99dB) | 0.77 (2.12dB) | 0.91 (1.63dB) |
| 38kHz | 0.96 (1.15dB) | 0.97 (0.78dB) | 0.93 (0.99dB) | 0.72 (1.37dB) | 0.55 (2.77dB) |
| 70kHz | 0.97 (0.91dB) | 0.93 (1.30dB) | 0.96 (0.68dB) | | |
| 120kHz | 0.97 (0.96dB) | 0.92 (1.36dB) | | | |
| 200kHz | 0.95 (1.21dB) | | | | |
| CCE1207 (https://doi.org/10.6075/J0TX3CPV) – $R^2$ (RMSE) | | | | | |
| 18kHz | | | | | |
| 38kHz | 0.96 (1.11dB) | 0.91 (1.91dB) | 0.86 (1.90dB) | 0.55 (2.16dB) | 0.43 (2.51dB) |
| 70kHz | 0.92 (1.64dB) | 0.92 (1.87dB) | 0.85 (1.73dB) | | |
| 120kHz | 0.96 (1.07dB) | 0.87 (2.17dB) | | | |
| 200kHz | 0.87 (1.67dB) | | | | |

| | 15-115m | 115-215m | 215-315m | 315-415m | 415-515m |
|---|---|---|---|---|---|
| NH0911 (https://doi.org/10.6075/J0T43RFB) – $R^2$ (RMSE) | | | | | |
| 18kHz | 0.93 (1.62dB) | 0.98 (0.89dB) | 0.98 (0.58dB) | 0.96 (0.71dB) | 0.89 (2.26dB) |
| 38kHz | 0.96 (0.98dB) | 0.97 (0.75dB) | 0.88 (1.58dB) | 0.62 (2.10dB) | 0.23 (3.12dB) |
| 70kHz | 0.96 (1.06dB) | 0.88 (2.12dB) | 0.38 (5.02dB) | | |
| 120kHz | 0.93 (1.38dB) | 0.58 (4.40dB) | | | |
| 200kHz | 0.69 (3.70dB) | | | | |
| MF1004 (https://doi.org/10.6075/J0JM27ZT) – $R^2$ (RMSE) | | | | | |
| 18kHz | 0.96 (1.83dB) | 0.98 (1.06dB) | 0.96 (0.87dB) | 0.96 (0.86dB) | 0.91 (1.90dB) |
| 38kHz | 0.95 (1.51dB) | 0.95 (1.28dB) | 0.97 (0.71dB) | 0.76 (1.06dB) | 0.83 (0.98dB) |
| 70kHz | | | | | |
| 120kHz | 0.96(1.34dB) | 0.91(1.76dB) | | | |
| 200kHz | 0.96 (1.41dB) | | | | |
| NH1011 (https://doi.org/10.6075/J0959FWD) – $R^2$ (RMSE) | | | | | |
| 18kHz | 0.96 (1.82dB) | 0.99 (0.83dB) | 0.98 (0.97dB) | 0.92 (1.07dB) | 0.90 (2.04dB) |
| 38kHz | 0.96 (1.05dB) | 0.99 (0.65dB) | 0.96 (1.74dB) | 0.83 (2.16dB) | 0.60 (2.93dB) |
| 70kHz | 0.98 (0.96dB) | 0.92 (2.95dB) | 0.57 (7.37dB) | | |
| 120kHz | 0.97 (1.04dB) | 0.71 (5.18dB) | | | |
| 200kHz | 0.96 (1.17dB) | | | | |
| CCE1106 (https://doi.org/10.6075/J0ZP44F1) – $R^2$ (RMSE) | | | | | |
| 18kHz | | | | | |
| 38kHz | 0.72 (2.01dB) | 0.87 (1.40dB) | 0.78 (1.06dB) | 0.82 (0.69dB) | 0.87 (0.95dB) |
| 70kHz | 0.06 (3.63dB) | 0.58 (3.10dB) | 0. (2.33dB) | | |
| 120kHz | 0.93 (0.94dB) | 0.85 (2.08dB) | | | |
| 200kHz | 0.82 (1.49dB) | | | | |
| NH1110 (https://doi.org/10.6075/J0NC5ZHB) – $R^2$ (RMSE) | | | | | |
| 18kHz | 0.98 (1.06dB) | 0.95 (1.92dB) | 0.85 (2.72dB) | 0.39 (4.03dB) | 0.40 (6.93dB) |
| 38kHz | 0.99 (0.77dB) | 0.60 (5.28dB) | 0. (9.74dB) | 0. (14.4dB) | 0. (13.7dB) |
| 70kHz | 0.98 (0.92dB) | 0.46 (3.62dB) | 0. (5.36dB) | | |
| 120kHz | 0.96 (1.23dB) | 0. (9.94dB) | | | |
| 200kHz | 0.81 (2.12dB) | | | | |
| OS1207 (https://doi.org/10.6075/J0862DTG) – $R^2$ (RMSE) | | | | | |
| 18kHz | | | | | |
| 38kHz | 0.97 (0.86dB) | 0.98 (1.06dB) | 0.95 (1.37dB) | 0.82 (1.59dB) | 0.90 (1.14dB) |
| 70kHz | 0.98 (0.69dB) | 0.98 (1.01dB) | 0.97 (1.00dB) | | |
| 120kHz | 0.97 (0.86dB) | 0.26 (9.79dB) | | | |
| 200kHz | 0.88 (1.58dB) | | | | |
| NH1210 (https://doi.org/10.6075/J04F1P35) – $R^2$ (RMSE) | | | | | |
| 18kHz | 0.96 (1.26dB) | 0.98 (1.13dB) | 0.98 (0.63dB) | 0.95 (0.62dB) | 0.83 (1.67dB) |
| 38kHz | 0.96 (0.90dB) | 0.97 (0.87dB) | 0.96 (0.92dB) | 0.88 (1.07dB) | 0.74 (1.44dB) |
| 70kHz | 0.97 (0.88dB) | 0.97 (0.91dB) | 0.96 (0.81dB) | | |
| 120kHz | 0.97 (0.98dB) | 0.87 (1.58dB) | | | |
| 200kHz | 0.94 (1.35dB) | | | | |

Figure 1: Performance of the acoustic data processing approach. For each transect, the nautical area scattering strength ($S_A$) is computed over 5 depth layers (15-115, 115-215, 215-315, 315-415, and 415-515 m), and averaged over 30 s intervals, for up to 5 frequencies (18, 38, 70, 120, and 200 kHz). Results from the processing approach used in this study are compared with the corresponding publicly available processed echograms. For each transect, we report the coefficient of determination ($R^2$) and the root mean square error ($RMSE$, in dB). At higher frequencies, only shallower layers are sampled due to stronger signal absorption (indicated by empty cells). Unavailable frequencies are shaded in dark gray. Echogram sections where the seafloor intersects with the depth layers are excluded.

### S3 Definition of functional groups

Multi-frequency acoustic signals provide coarse information on the nature of the organisms sampled. The Multi-Frequency Indice (MFI) [1] calculated from measured volume backscattering coefficients ($s_v$), at 18, 38, 70, 120 and 200 kHz, provide a coarse characterization of the dominant organism groups in acoustic observations, including fish with swimbladder (MFI<0.4) and fluid-like zooplankton (0.7<MFI<0.8).

In our analysis, 28 cruises out of 41 sampled each of the five frequencies required to compute MFIs, while all sampled at least 38 and 120 kHz, two frequencies from which the difference $\Delta S_{v\ 120-38\,kHz} = S_{v\ 120\,kHz} - S_{v\ 38\,kHz}$ can be used to coarsely determine dominant functional groups sampled [2, 3]. To use all available cruises, we assign categories of swimbladdered 'fish' and 'zooplankton' based on $\Delta S_{v\ 120-38\,kHz}$ determined from the ranges of values for MFI<0.4 or 0.7<MFI<0.8 in the subset of cruises with all five frequencies (see Fig. 2).

In the upper ocean layer, [15-175] m, the range of $\Delta S_{v\ 120-38\,kHz}$ values that include 95% of swimbladdered fish (MFI<0.4) is $-9$ to 0 $dB\ re\ 1\ m^{-1}$; 95% of zooplankton (0.7<MFI<0.8) occur within the range 0 to 16.5 $dB\ re\ 1\ m^{-1}$. Observations with $\Delta S_{v\ 120-38\,kHz}$ $-9 <$ or $>$ 16.5 are labeled as 'other' and not discussed in the analysis.

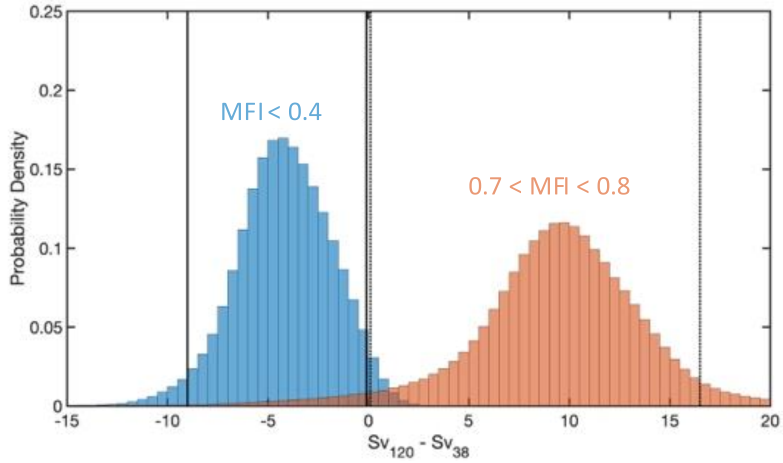


Figure 2: Probability density distribution of $\Delta S_{v\ 120-38\,kHz}$ for swimbladdered fish (MFI<0.4, in blue) and zooplankton (0.7<MFI<0.8, in orange) based on echograms that include 18, 38, 70, 120 and 200 kHz frequencies. The plain and dotted vertical lines show the approximate 5th and 95th percentiles for each distribution.

### S4 Results based on calibrated transects

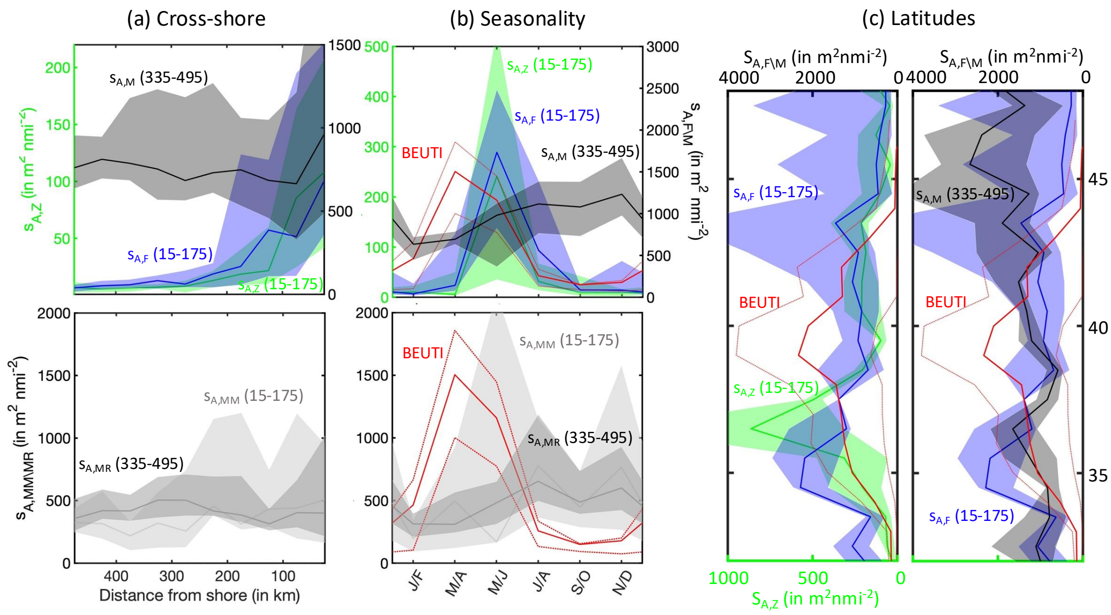


Figure 3: Backscatter variability from epipelagic to mesopelagic layers based on calibrated data only, (a) cross-shore, (b) seasonal and (c) latitudinal distributions. Panels show nautical area scattering coefficient (NASC, or $s_A$, in $m^2\ nmi^{-2}$) for surface epipelagic zooplankton $s_{A,Z}$ (green), fish $s_{A,F}$ (blue), mesopelagic fish (335-495 m) $s_{A,M}$ (black), and migrating mesopelagic $s_{A,MM}$ (light grey) or resident mesopelagic fish $s_{A,MR}$ (dark grey). Solid lines show the median across 11 years of observations, and envelopes the 25th-75th percentiles. The solid red line shows the mean BEUTI index–the dotted lines the 25th-75th percentiles across years–in Southern California [4].

### S5 Cross-shore driver distributions

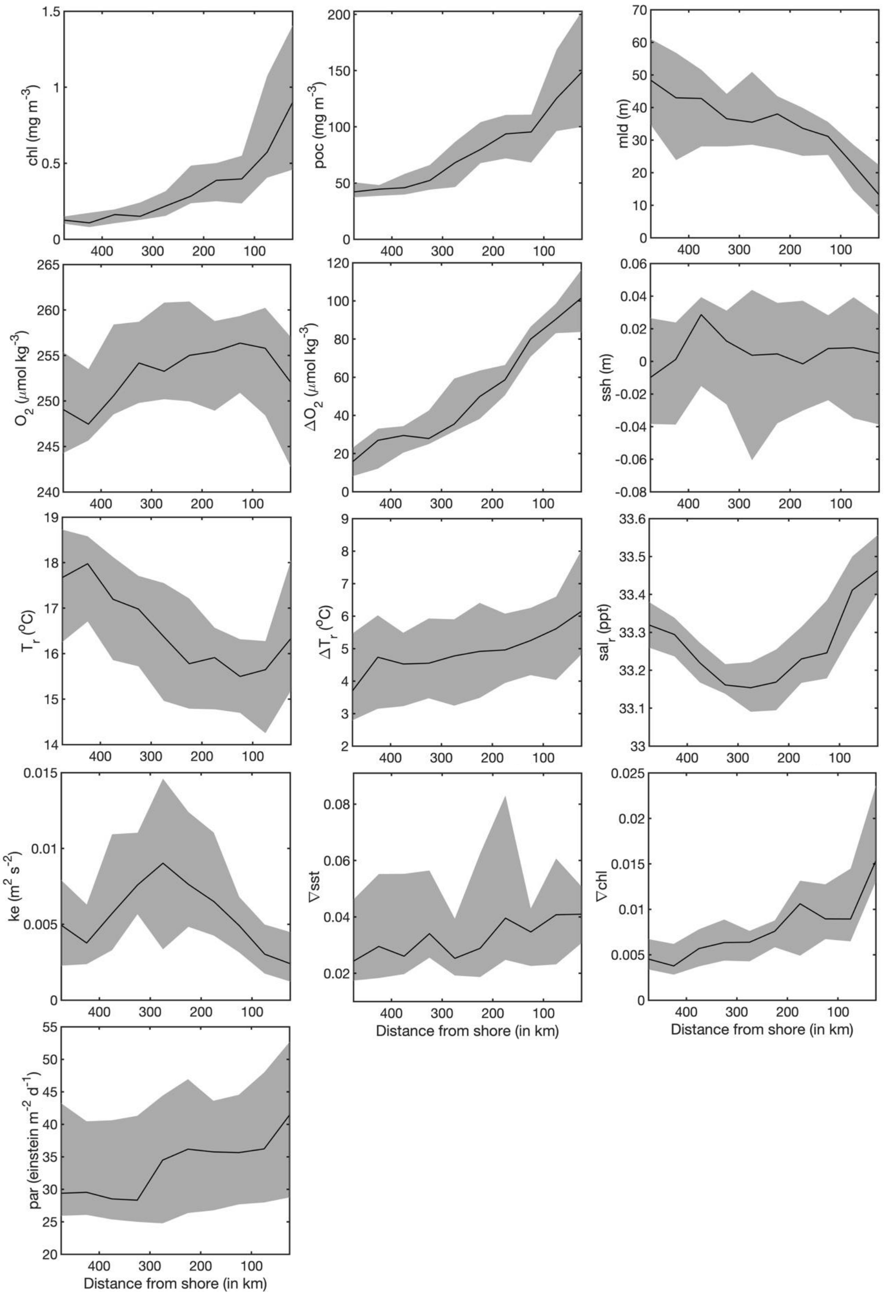


Figure 4: Environmental drivers along a cross-shore gradient. The solid line represents the median observations across 11 years and the envelope the 25th-75th percentiles.

### S6 Seasonal driver distributions

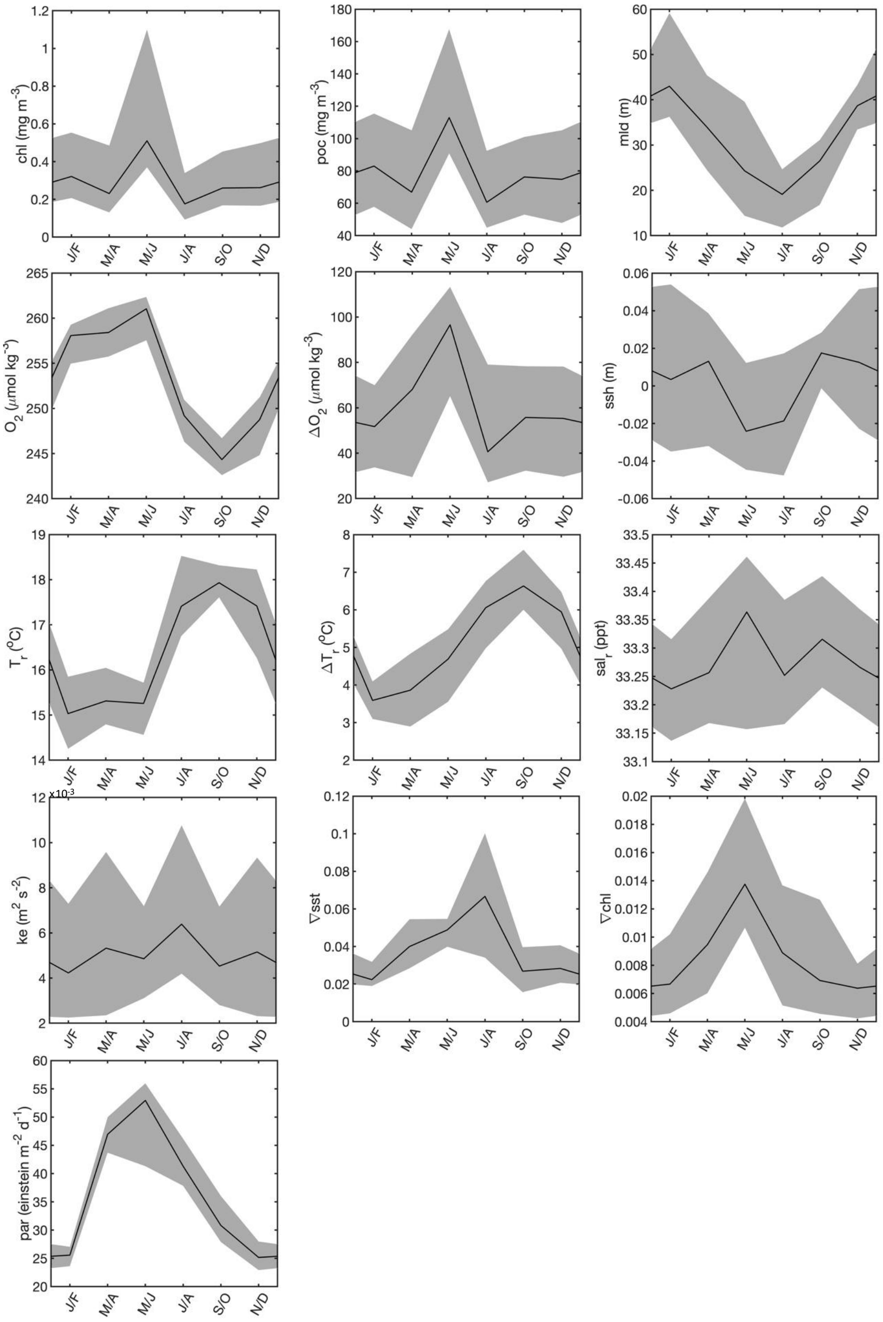


Figure 5: Seasonal cycle of environmental drivers. The solid line represents the median observations across 11 years and the envelope the 25th-75th percentiles.

### S7 Latitudinal driver distributions

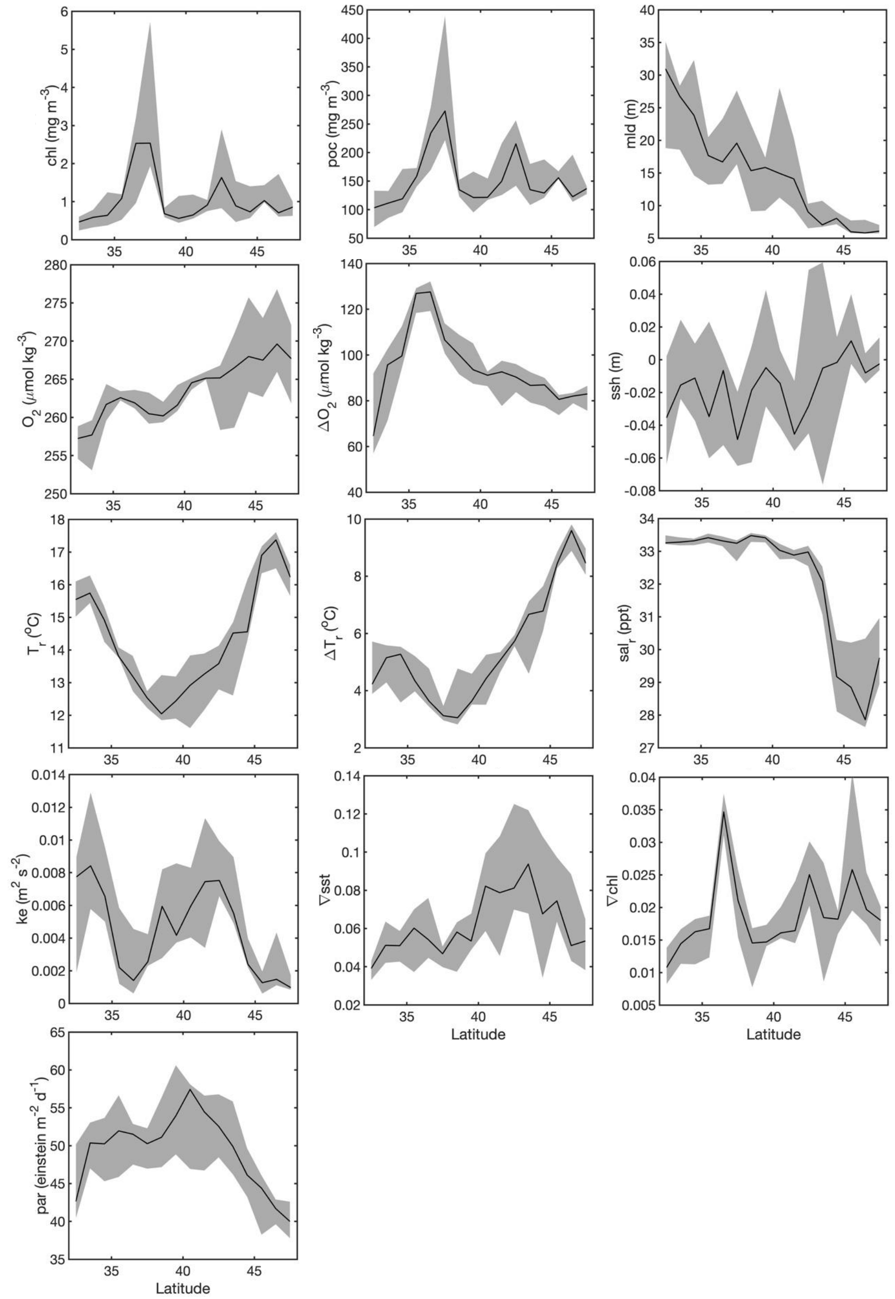


Figure 6: Environmental drivers along a latitudinal gradient. The solid line represents the median observations across 11 years and the envelope the 25th-75th percentiles.

### S8 Environmental correlations for indicators of vertical distribution

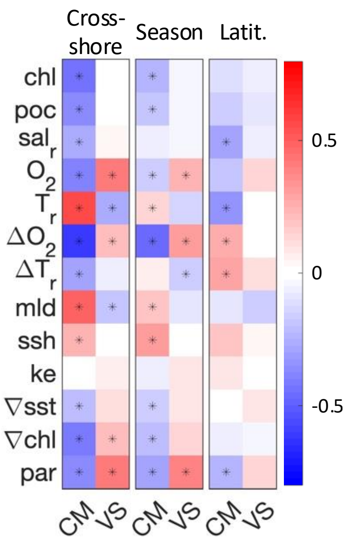


Figure 7: Correlation between environmental variables and indicators of vertical MTL distribution—center of mass (CM) or vertical spread (VS)—based on linear models where cross-shore, seasonal, or alongshore variations are treated as a fixed effect. Asterisks (∗) denote correlations with p-value $< 0.001$.
